# Febrile temperature activates the innate immune response by promoting aberrant influenza A virus RNA synthesis

**DOI:** 10.1101/2025.05.19.654939

**Authors:** Karishma Bisht, Daniel R. Weilandt, Caitlin H. Lamb, Elizaveta Elshina, Cameron Myhrvold, Aartjan J.W. te Velthuis

## Abstract

Fever during influenza A virus (IAV) infection is triggered by the innate immune response. Various factors contribute to this response, including IAV mini viral RNAs (mvRNA), which trigger RIG-I signaling when their replication and transcription are dysregulated by template loops (t-loop). It is presently not well understood whether the fever response to IAV infection impacts subsequent viral replication and innate immune activation. Here we show that IAV infection at temperatures that simulate fever leads to increased antiviral signaling in H1N1 and H3N2 infections. Mathematical modeling and experimental analyses reveal that differential IAV nucleoprotein and RNA polymerase production increase mvRNA and interferon production. Moreover, at the higher infection temperature, mvRNAs with dysregulating t-loops contribute most to the innate immune activation. We propose that fever during IAV infection can establish a positive feedback loop in which elevated aberrant RNA synthesis and innate immune activation can contribute to the dysregulation of cytokine production.

## Introduction

Influenza A viruses (IAV) cause moderate to severe respiratory disease in seasonal epidemics and occasional pandemics^1^. The level of activation and dysregulation of the innate immune response contributes to the outcome of IAV infection, particularly in infections with highly pathogenic avian or pandemic IAV strains^2,3^. Activation of the innate immune response involves the binding of IAV RNA to host pathogen receptor retinoic acid-inducible gene I (RIG-I)^4–6^. Once activated, RIG-I initiates the oligomerization of MAVS, triggering the phosphorylation of NF-kB, IRF-3 or IRF-7 and the subsequent expression of interferons (IFN) type I and III and proinflammatory genes, including tumor necrosis factor (TNF) and interleukin 6 (IL6)^7,8^. In turn, IFNs drive STAT1 and 2 phosphorylation, and the expression of IFN-stimulated genes (ISG). Proinflammatory genes and ISGs have many roles in IAV infection, including recruitment of immune cells, activation of negative feedback loops, and triggering the febrile response that elevates the host temperature from approximately 37°C to 38.5-41.0°C^9,10^. While fever has been known to be a common and early IAV infection symptom for decades, its impact on viral replication and gene expression is understudied.

The IAV genome consists of eight segments of single-stranded, negative sense viral RNA (vRNA) that exist in the context of viral ribonucleoproteins (vRNPs) and are bound by at least 20 NP^11,12^. The replication and transcription of the viral genome are complex processes that involve both viral and host factors (Fig. 1a). The IAV RNA polymerase first produces capped and polyadenylated mRNA molecules during IAV transcription. Next, the IAV RNA polymerase produces a complementary RNA (cRNA) that is subsequently copied into a new vRNA molecule as part of viral replication. In addition, the IAV RNA polymerase produces various aberrant or non-canonical RNA molecules, including deletion-containing viral genomes (DelVGs), mini viral RNAs (mvRNA), and capped cRNAs (ccRNA) (Fig. 1a)^13–17^. DelVGs and mvRNAs are diverse in sequence, lack internal gene sequences, but retain the conserved the 5’ and 3’ termini of full-length vRNA segments that act as viral promoter. DelVGs and mvRNAs can therefore be replicated and transcribed. The difference between the two molecules is that mvRNAs are sufficiently short (<125 nt) to be replicated and transcribed in the absence of viral nucleoprotein (NP) and that a reduction in NP levels stimulates mvRNA formation. In contrast to mvRNAs and DelVGs, ccRNAs are IAV transcripts that contain a cap, but lack a polyA-tail (Fig. 1a)^15,16^. Presently the role of mvRNAs and ccRNAs is not fully understood. One role of mvRNAs and ccRNAs in infection is activation of RIG-I signaling. mvRNAs that contain a so-called template loop (t-loop) contribute most to RIG-I activation, because the t-loop causes ccRNA formation during mvRNA transcription and RNA polymerase stalling during mvRNA replication, which together lead to the release of mvRNAs and the putative formation of mvRNA-ccRNA duplexes^15^. While IAV replication and transcription have been captured with computational models previously, they have not been combined into a single model that also captures IAV aberrant RNA synthesis and innate immune signaling.

**Fig. 1:**
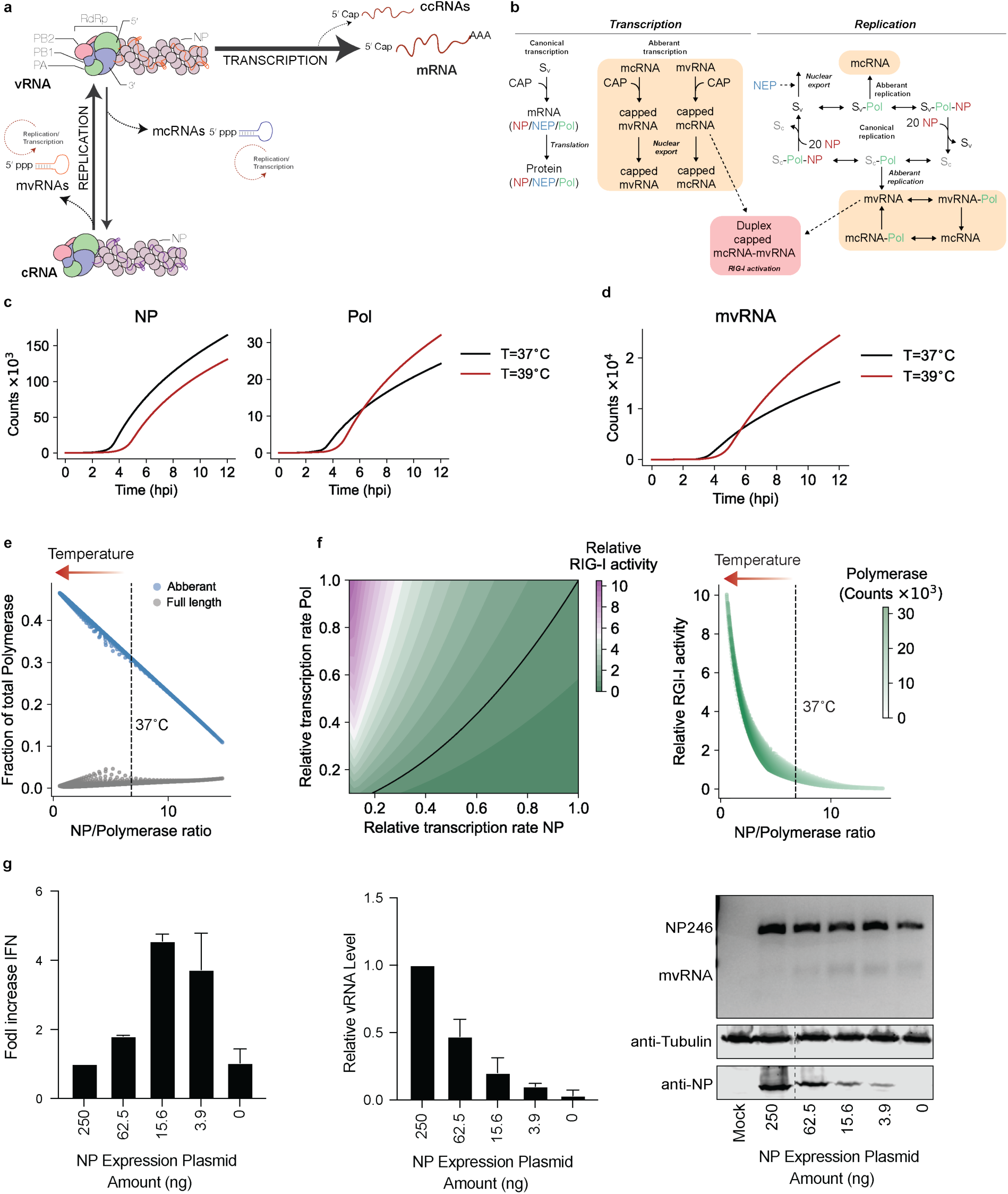
The single infection cycle model of influenza A Virus (IAV) reveals impact of NP dynamics on innate immune activation. **a,** Schematic of IAV replication and transcription during IAV infection. The viral ribonucleoprotein (vRNP) consists of a copy of the viral RNA-dependent RNA polymerase (RdRp), vRNA (orange), and NP (purple). The RdRp consists of three subunits: PB1 (blue), PA (green) and PB2 (pink). Canonical transcription of vRNA molecules produces mRNAs, while replication of vRNA produces cRNAs (purple). cRNA molecules are encapsidated into cRNPs. Non-canonical transcription of vRNAs can produce ccRNAs, while non-canonical replication of vRNA molecules produce mcRNAs. Also, cRNA molecules can be replicated into mvRNAs. The mcRNAs and mvRNAs can be transcribed and replicated (dotted circles). **b,** Schematic illustrating the processes and variables (see Table S2) incorporated into the model. **c,** Simulation of IAV infection cycle and prediction of NP and polymerase protein levels per cell. **d,** Simulation of IAV infection cycle and prediction of mvRNA levels per cell. **e,** Replication rate of aberrant and full-length products driven by the NP/polymerase ratio. **f,** Heatmap on left shows RIG-I activity as function of NP and polymerase gene transcription. Plot on right shows RIG-I activity as function of the ratio between the NP and polymerase proteins that follows from the simulations in left plot. Arrow indicates shift in RIG-I activation after fever. **g,** Analysis of *IFNB* promoter activity and vRNA levels using primer extension during the replication of a segment 5-based 246-nt RNA template by the WSN polymerases. NP mvRNA levels were analyzed by RT-PCR. NP expression was assessed by western blot (bottom gel). *n* = 3 biologically independent experiments.

IAVs can infect cells of the upper and lower respiratory tract, and the viral replication machinery may thus experience different temperature environments during infection^18–20^. In addition, the IAV replication machinery may experience a change in temperature when innate immune signaling activates the febrile response. Previous in vitro studies have shown that IAV RNA synthesis is altered when the incubation temperature is increased^21^, but the impact of the temperature on IAV infection in cells is still poorly understood. A potential confounding factor in such a study is that simulation of the febrile response through heat-shock will affect host cell gene expression, host post-translational modifications, as well as viral enzymatic activity. These combinatorial effects make it difficult to measure how an increase in temperature impacts the efficiency of IAV replication and transcription, and how putative aberrant RNAs produced under this condition affect the innate immune system. Studying the impact of temperature on IAV infection in a mouse model is also complicated, as the mice either enter hypothermia or show no fever response following IAV infection^22^.

To explore the impact of temperature on the multiple viral processes as well as study the interactions among IAV replication, transcription, aberrant RNA synthesis and innate immune activation, we used mathematical modeling to simulate a single infection cycle. Our modeling predicted that a change in infection temperature from 37°C to 39°C would lead to dysregulated viral nucleoprotein (NP) and polymerase acidic (PA) subunit levels, and an increase in mvRNA production and innate immune signaling. This change in temperature simulates a mild fever response, similar to what is seen with the flu symptoms observed from both seasonal and pandemic strains. We subsequently designed experiments to verify our model and measure the effect of temperature on IAV RNA synthesis. Specifically, we adapted cells to 37 or 39°C prior to infection or transfection to minimize the impact of heat-shock and avoid differential gene expression between cells growing at either temperature. We find that our experimental results align with our model and demonstrate that a febrile-like temperature enhances mvRNA synthesis and activation of the RIG-I-dependent innate immune response. We also find that activation of RIG-I is dependent on the mvRNA sequence as well as mvRNA transcription, in line with recent results showing that IAV transcription contributes to innate immune activation^15^. When we compare pandemic and adapted IAV RNA polymerases, we find that the transcription of mvRNAs by a pandemic IAV RNA polymerase is significantly increased at 39°C. Finally, we performed infection experiments to show that the temperature-dependent modulation of the innate immune response is preserved across infections with both H1N1 and H3N2 influenza A virus strains. Overall, we speculate that, depending on the IAV strain and base level of mvRNA synthesis, activation of the febrile response could create a positive feedback loop that could contribute to dysregulation of the immune system and production of a cytokine storm.

## Results

### Modeling of influenza virus replication and transcription dynamics

mvRNA and full-length vRNA segment replication and transcription likely occur simultaneously as an IAV infection progresses. Since these processes may affect each other as well as the accumulation of viral proteins, innate immune signaling, and the biochemical activity of the viral replication complex, it is hard to intuit how changes in this network of viral and host interactions affect the outcome of an infection. To better understand how a change in temperature impacts the dynamics of this network, we developed a mathematical model for a single IAV replication cycle. Building on previous IAV infection models^23^, we account for nuclear export, the formation of mvRNAs, activation of RIG-I, and explicitly define IAV cap-snatching of host (pre-)mRNAs caps as a limiting resource (and thus that other resources like NTPs, lipids, and amino acids are in excess). Additionally, we incorporated recent biochemical and structural findings, including that replication and transcription occur in a complex consisting of the viral RNA polymerase and host factor ANP32 (we define this as a single complex Sv), that NP is required for full-length segment replication, but not mvRNAs, and that mvRNAs can be replicated and transcribed (Fig. 1b). In addition, we assumed that mvRNAs and their transcription products (in particular ccRNAs) can be exported from the nucleus and bind RIG-I to activate an innate immune response^15^, that the replication of the eight IAV segments (producing vRNA and cRNA) is not fundamentally different, and that expression differences among the segments are driven by variations in transcription (mRNA) and/or translation efficiency. Finally, we simplified the existing viral infection model by accounting for only three protein complexes: i) the viral RNA polymerase subunit PA bound to PB1, PB2 and ANP32, ii) the viral NP, and iii) the viral nuclear export protein (NEP) bound to M1 and CRM1. For simplicity and to ensure that experimentally testing the model remained feasible, we assumed that other viral proteins, including PA-X, PB1-F2, PB1-N40, NS1, HA, NA, and M2, were not altered by the change in temperature. We parametrized the model using previously published parameters and added biophysical estimates for the added processes (Table S1^24–29^). The resulting model reasonably predicts the abundance of the canonical viral RNA species during a single infection cycle in comparison to experimental data (Fig. S1a-d)^30^.

### Modeling predicts that an increase in temperature lowers NP expression and increases mvRNA expression and innate immune signaling

An increase in temperature above physiological conditions reduces the binding affinities of protein-ligand interactions and the catalytic function of enzymes. Previous experimental results have shown that the binding affinity of the IAV RNA polymerase to RNA is indeed reduced at higher temperatures^21^. However, simulation with reduced binding affinities showed that this change does not inherently increase the rate of mvRNA formation (Fig. S1e, f). Instead, simulating IAV replication at 37°C and 39°C predicts that NP levels become reduced from ∼2 hpi at 39°C, while the polymerase levels become elevated from ∼5 hpi (Fig. 1c). Decreasing the NP level results in reduced processivity or a failure to encapsidate full-length viral segments into RNPs, and our model predicts that mvRNAs levels at 39°C start to outpace mvRNAs levels at 37°C from 5 hpi onwards at the cost of full-length RNA synthesis (Fig. 1d, e). The imbalance between NP and the viral RNA polymerase also accelerates the accumulation and transcription of mvRNAs in the cell and the potential of mvRNA-ccRNAs to activate RIG-I (Fig. S1g). Thus, the imbalance in the NP to viral RNA polymerase ratio (NP:polymerase <7) impacts RIG-I activation (Fig. 1f). The effect of this dynamic can be shown experimentally when we titrate NP in a mini-genome assay, using a 246-nt long template from the NP-encoding segment (NP246): as NP availability to the RNA polymerase decreases, full-length template RNA levels decrease, while mvRNA levels and IFN-beta promoter activity increase (Fig 1g), in line with previous observations^14^. Based on these findings, we propose that the activation of RIG-I is increased at 39°C and particularly sensitive to decreased nuclear protein expression at this temperature.

### Temperature elevation upregulates the expression of innate immune genes during IAV infection

To confirm the predictions of our model experimentally, we performed A/WSN/33 H1N1 (WSN) infections at 37 or 39°C in temperature-adapted A549 cells at MOI of 3. We extracted total RNA at 12 hpi, quantified host and viral gene expression using RNAseq (Fig. 2a), and used edgeR to identify differentially expressed genes (DEGs). The only DEGs between the mock infected cells adapted to 37 or 39 °C were 3 non-innate immune genes (Fig. S2a). We observed a comparable result in temperature-adapted HEK293T cells, in which only a few non-innate immune genes were DEGs (Fig. S2b). However, in the infected A549 cells, we found that 31 genes were differentially upregulated at 39°C relative to 37°C (Fig. 2b). The DEGs included known innate immune signaling factors ISG1, RIG-I, and IRF7 (Fig. 2b). A reactome pathway enrichment analysis confirmed that DEGs at 39 compared to 37°C were notably enriched in type I interferon, antiviral defense, innate immunity, and ISG15 signaling pathways (P < 0.05, FDR < 0.05) (Fig. 2c). Together, these results indicate that IAV infection at a simulated febrile temperature causes an upregulation in antiviral gene expression.

**Fig. 2:**
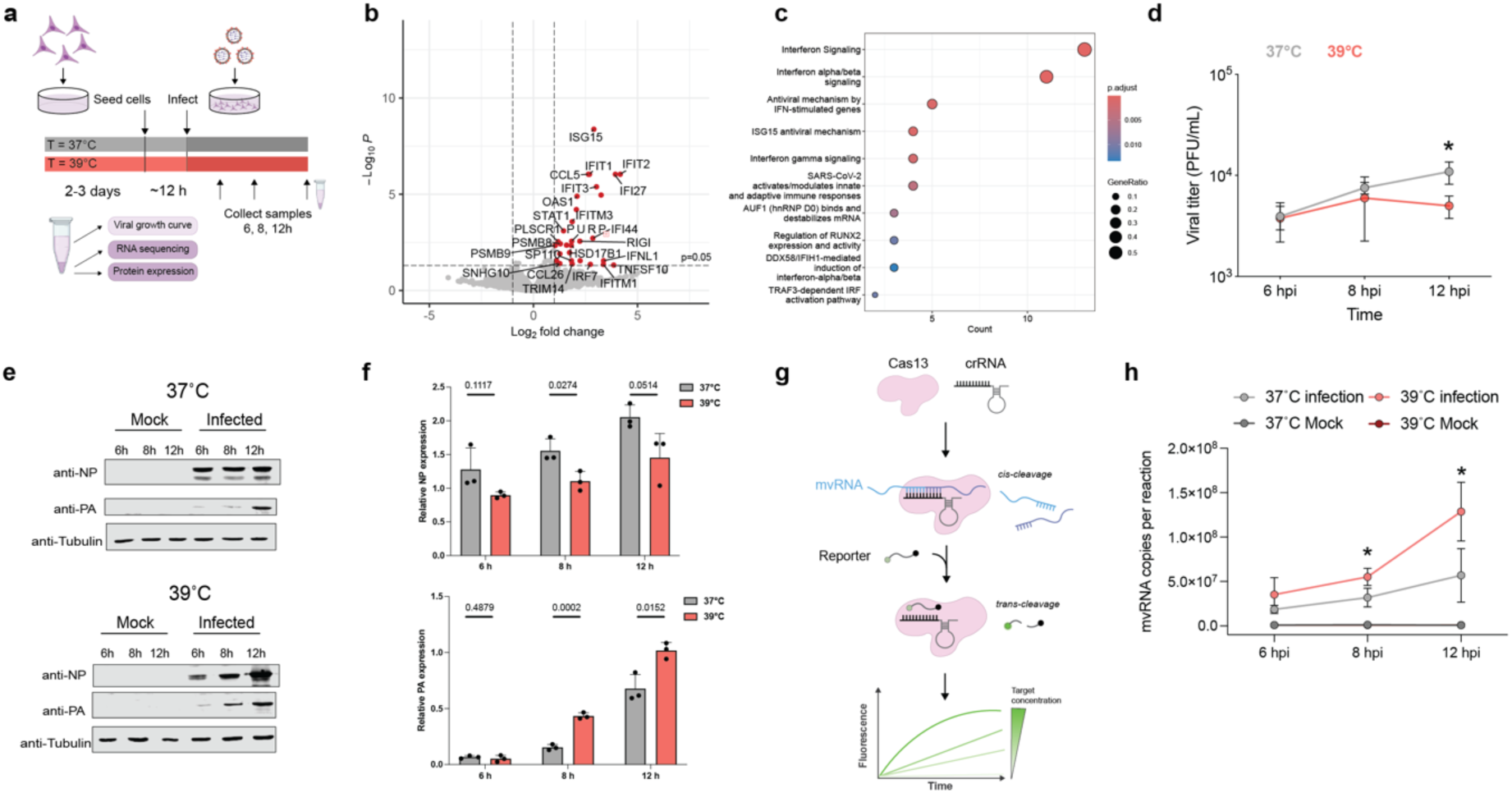
Elevated temperature affects both innate immune response and aberrant RNA synthesis during infection. **a,** Schematic of experimental design used for viral infection of A549 cells in cells adapted at 37°C and 39°C. **b,** Volcano plot illustrating differentially expressed genes in A549 cells infected with A/WSN/33 virus at an MOI of 3. RNA-seq was performed on 3 biological replicates. **c,** Reactome pathway enrichment analysis of the differentially expressed genes. **d,** Growth kinetics of lab adapted WSN virus in A549 cells at different temperatures (37°C, or 39°C). A549 cells were infected at an MOI of 3 at 37°C and 39°C. The supernatants of the infected cells were harvested at the indicated times, and the virus titers were determined by performing plaque assays in MDCK cells at 37°C. **e,f,** Viral protein expression analyzed using western blot, the bar graph depicts the quantified relative expression level. **g,** Schematic of RNA detection using Cas13. **h,** Detection of NP-61 mvRNA using Cas13 assay in fractionated A549 cells infected with WSN. Graph shows the copy number of mvRNA NP-61 in infected cells at 37°C and 39°C. Data are shown as a mean of 3 independent experiments. Error bars indicate standard deviation. The *P* values were determined by using an unpaired *t* test. (\**P* < 0.05).

### Temperature elevation increases synthesis of non-canonical RNA products

The transcriptomics data showed a differential response to the IAV infection at 39°C that was independent of the effect of temperature on the transcriptional landscape of the cell (Fig. 2b-c). To understand how IAV increases innate immune activation in a temperature-dependent manner, we first measured virus growth. As shown in Fig. 2d and Fig. S3a-b, we observed that IAV titers were modestly, but significantly reduced at 39°C relative to 37°C in both A549 and HEK293T cell lines, in line with other reports^31^. Western blot analysis ruled out that heat-shock protein 70 (hsp70), which is known to inhibit influenza virus replication, was responsible for the lower titer levels observed at 39°C (Fig. S3d). Analysis of the viral PA and NP levels showed a reduction in NP expression and an increase in PA expression at 39°C, both in a high and low MOI infection setting (Fig. 2e-f, Fig. S3c), suggesting that either viral mRNA levels or viral protein synthesis were differentially affected for these two viral genes at this temperature. These results were in line with the effect of temperature predicted by our model (Fig. 1).

We next investigated if mvRNA synthesis was increased during infection at 39°C relative to 37°C. Using an amplification-free CRISPR-Cas13 assay that targets the unique mvRNA junction sequence (Fig. 2g), thereby distinguishing mvRNAs from full-length NP-encoding vRNAs, we quantified a 61 nt-long NP-encoding segment derived mvRNA (NP-61). This mvRNA is highly abundant in ferret lung tissue infected with A/Brevig Mission/1/1918 (H1N1) or A/Indonesia/2005 (H5N1) and A549 cells infected with WSN^14,32,33^. As shown in Fig. 2h, we observed a significant increase in the copy number of the NP-61 mvRNA in cells infected with IAV at 39°C relative to 37°C. These results suggest that mvRNAs are produced at higher levels at 39°C compared to 37°C.

### Temperature influences NP and polymerase proteins in distinct ways during an IAV infection

To further validate the predictions of our model and investigate the mechanism underlying the differential NP and PA expression, we infected A549 cells at an MOI of 1 and examined viral RNA and NP and PA protein levels (Fig. 3). Quantification of the segment 3 and 5 vRNA and capped RNA (mRNA and ccRNA) levels by primer extension revealed an increase over the course of the infection, with a more pronounced increase at 39°C relative to 37°C (Fig. 3a-d, Fig S4a-d). Western blot analysis (Fig. 3e-f) showed opposite trends for the NP and PA protein levels: NP was more highly expressed at 37°C compared to 39°C, while the PA RNA polymerase subunit was more highly expressed at 39°C. These findings align with our MOI 3 experiments (Fig. 2) and they are in line with the decrease in NP expression previously observed in cells infected with IAV and IBV when incubated at 39°C^31^.

**Fig. 3:**
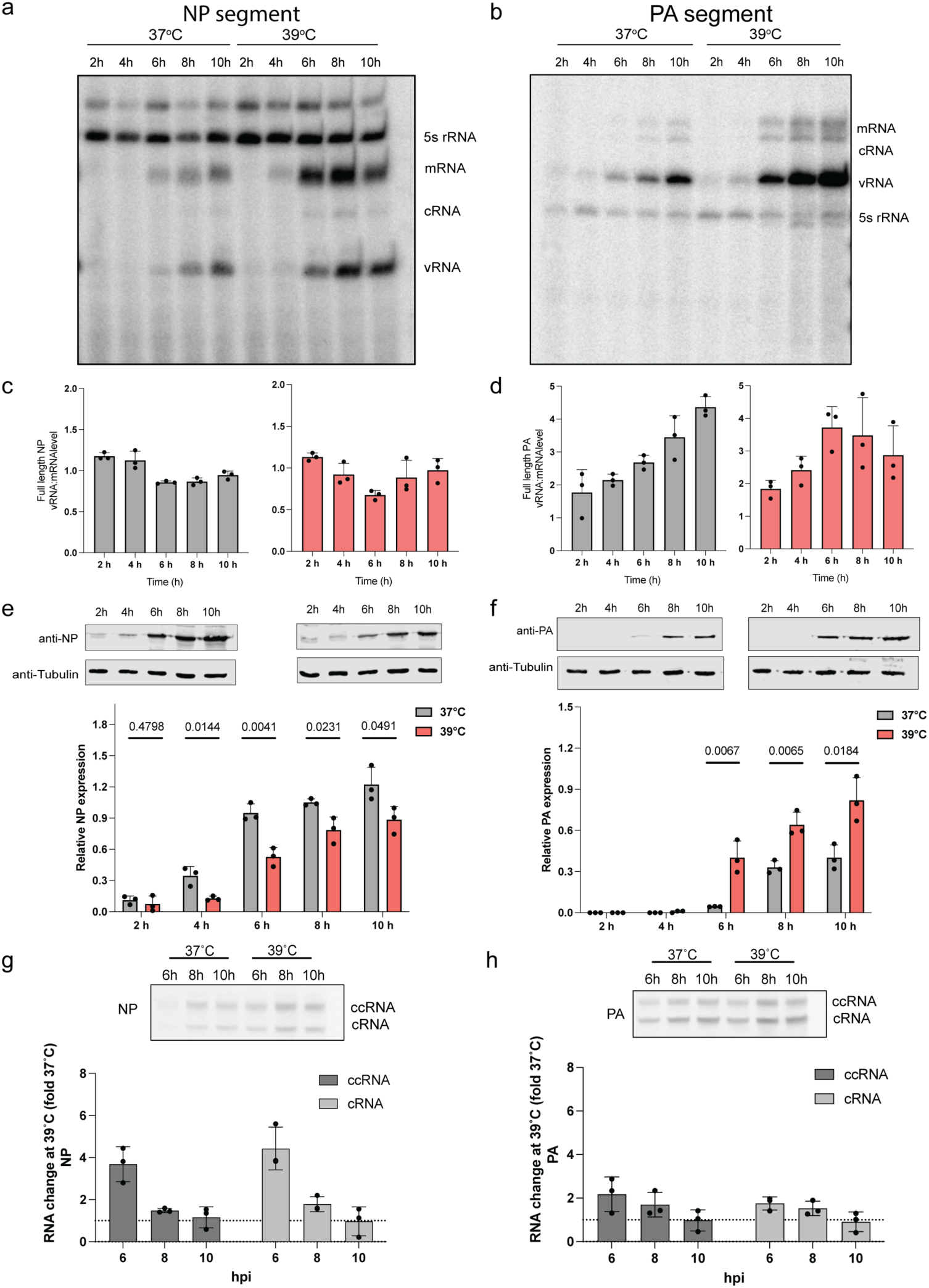
Differential NP and polymerase protein expression at elevated temperature during infection. A549 cells adapted at 37°C and 39°C were infected with an MOI of 1 of A/WSN/33 virus and RNA samples were taken at different time points post-infection. **c,d,** Normalization of vRNA levels relative to mRNA levels for the NP and PA full length segments. **e,f,** Cell lysates were collected at different time points and analyzed by immunoblotting. Lower panel show the quantified western blot data. **g,h,** Detection of ccRNA and cRNA by TSO-based RT-PCR. Quantification of fold-change difference in ccRNA and cRNA levels produced at 39°C versus 37°C. Data are shown as the mean of 3 independent experiments. Error bars indicate the standard deviation. The *P* values were determined by using an unpaired *t* test.

Our results consistently indicate that NP-encoding mRNA levels are high, but that NP protein levels are reduced at 39°C (note that this is below the melting temperature of NP, which is ∼45°C^34^). We hypothesized that this discrepancy between the IAV mRNA and protein level might be due to non-canonical transcription termination, which can result in the formation of ccRNAs. In primer extension experiments (Fig. S4c-d), ccRNA and mRNA molecules are indistinguishable from each other, but ccRNAs can be measured using an RT-PCR that is specific for the 3’ terminal sequence^35^. Following mRNA depletion and RT-PCR detection, we found an ∼4-fold increase in the NP-encoding segment ccRNA levels at 39°C compared to 37°C early in infection, while for PA-encoding segment, we observed only a 2-fold increase in the ccRNA level at 39°C. Together, these data suggest that segment 5 transcription is more strongly affected by non-canonical transcription termination than segment 3 transcription (Fig. 3g-h) and that the ccRNA ratio for segment 5 is almost 2-fold higher than segment 3. Given that it is currently unclear whether ccRNAs can be translated into protein, we propose that the observed increase in ccRNA levels might contribute to the decreased NP protein expression at 39°C.

### Temperature affects mvRNA replication and transcription in a t-loop dependent manner

mvRNAs are diverse in sequence^33^, but only mvRNAs that contain a t-loop can activate RIG-I^15^. We previously showed that the RNA polymerases from both human and avian-adapted IAV strains are sensitive to t-loops. mvRNAs can only be studied using transfection experiments, since currently no tools exist to manipulate individual mvRNAs during infection. To confirm that the t-loop-containing mvRNAs also triggered higher IFN-β reporter activities at 39°C, we transfected the IAV RNA polymerases from WSN or A/Brevig Mission/1/18 H1N1 (BM18) alongside an IFN-β reporter and one of six mvRNAs derived from three different segments: PA-60 and PA-66, which are derived from segment 3, NP-71.11 and NP-71.1, which are derived from segment 5, and PB1-62 and PB1-66, which are derived from segment 2. For the transfection, we used 39°C-adapted HEK293T cells, which show no difference in innate immune signaling compared to cells adapted to 37°C (Fig. S2b). Luciferase signals were measured twenty-four hours after transfection and were normalized to the co-transfected *Renilla* luciferase transfection control. Our mini genome assay revealed that mvRNAs that contained a t-loop (NP-71.11, PA-66, and PB1-66) induced a significantly higher IFN-β reporter activity at 39°C than 37°C (Fig. 4a-c, Fig. S5a-c). In contrast, the IFN-β reporter activity between the two temperatures was not different for mvRNAs that did not contain a t-loop (NP-71.1, PA-60, PB1-62) (Fig. 4a-c, Fig. S5a-c). We observed these differences in IFN-β reporter activity for both the WSN and BM18 RNA polymerases. As a control, we compared the IFN-β reporter activity induced by the replication of full-length segments 2, 3 and 5, and observed no significant difference between the two temperatures (Fig. S9a). Together, these results indicate that an increased activation of the innate immune response at 39°C is dependent on how transient RNA structures modulate IAV RNA polymerase activity on mvRNAs.

**Fig.4:**
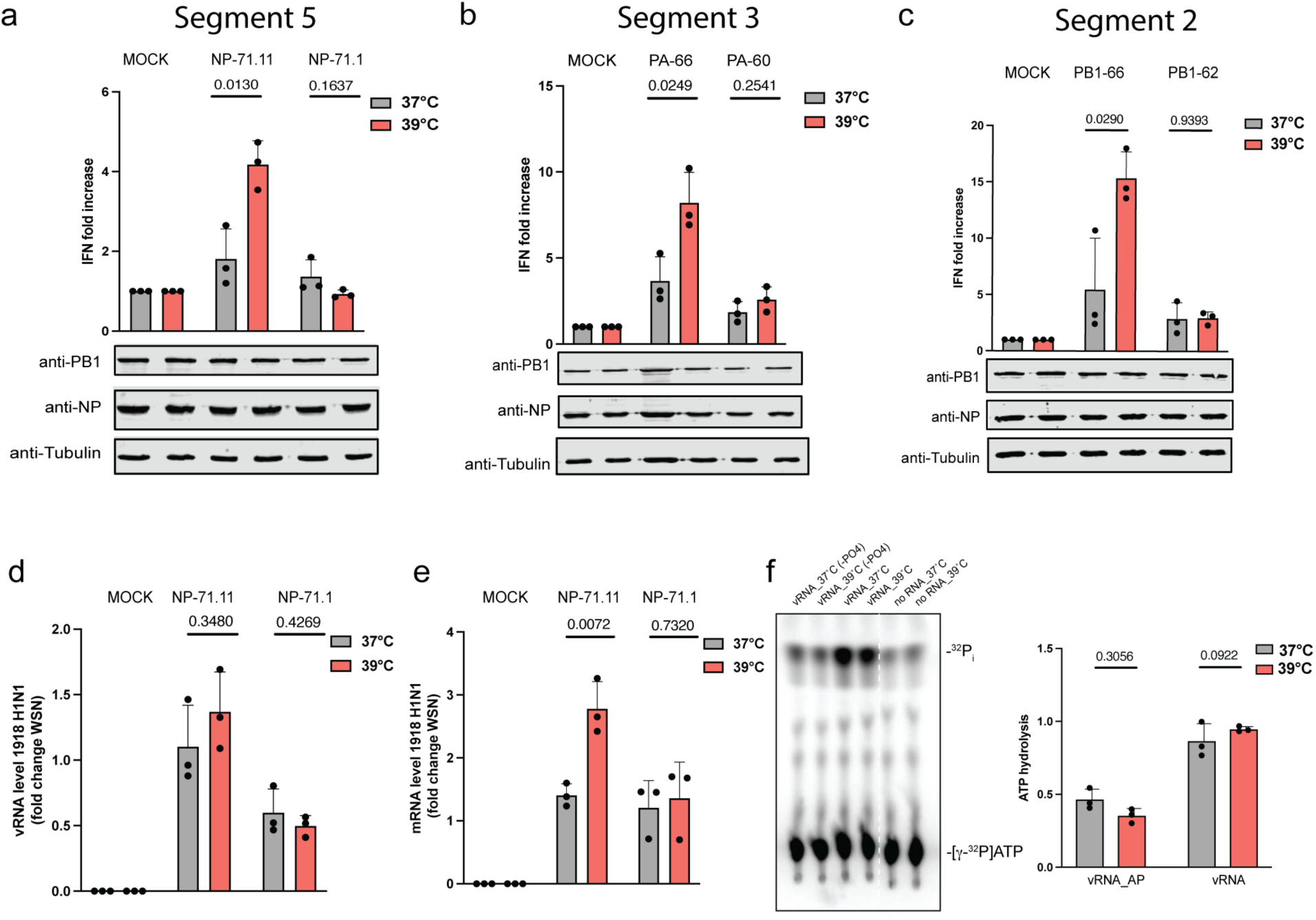
mvRNAs induce increased innate immune signaling at higher temperature. **a,b,c,** Analysis of IFN-β promoter activity induced by the replication of segment 5, 3 or 2 mvRNAs 24 hours post-transfection of BM18 IAV RNA polymerase. PB1, NP and tubulin expression was analyzed by western blot. **d,** Normalized vRNA levels for BM18 polymerase relative to WSN polymerase. **e,** Normalized mRNA levels for BM18 polymerase relative to WSN polymerase. **f**, ATPase activity of recombinant RIG-I was assessed in the presence of *in vitro* transcribed PA66 mvRNA, dephosphorylated PA66 mvRNA, or a no RNA control. Data are shown as the mean of 3 independent experiments. Error bars indicate standard deviation. *P* values were determined using a two-sided, unpaired *t*-test.

We previously showed that t-loops affect IAV RNA polymerase processivity^33^. To investigate whether the increased IFN-β reporter activity was caused by a further decrease in RNA polymerase activity, we extracted RNA from the mini-genome assay and measured viral RNA levels using primer extension analysis. For the BM18 RNA polymerase, two of the mvRNAs with a t-loop (NP-71.11 and PA-66) showed reduced vRNA steady-state levels at 39°C compared to 37°C (Fig. S6a-b), while the third with a t-loop, PB1-66, did not show a significant reduction in vRNA steady state level between the two temperatures (Fig. S6c). The three mvRNAs without a t-loop (NP-71.1, PA-60, and PB1-62) showed no difference in the vRNA steady-state levels between the two temperatures (Fig. S6a-c). For the WSN RNA polymerase, we observed reduced replication steady-state levels for all three mvRNAs with a t-loop and no differences for the three mvRNAs without a t-loop (Fig. S7a-c).

For the mRNA steady state levels produced by the BM18 RNA polymerase, we found no significant differences for all six mvRNA templates, with or without the t-loop, between 37°C and 39°C (Fig. S6d-f). However, in the presence of the WSN polymerase, mvRNAs NP-71.11 and PA-66 showed a notable decrease in mRNA levels, while the other four templates showed no changes (Fig. S7d-f). Finally, we measured the production of ccRNA molecules and observed a significant decrease for NP-71.11 and PA-66 templates with both the BM18 H1N1 and WSN polymerases at 39°C compared to 37°C. For the other four mvRNA templates, we observed no reduction in ccRNA level (Fig. S6g-i, Fig. S7g-i). Given that there were slight differences between the steady state RNA levels produced by the BM18 and WSN RNA polymerase, we compared the relative vRNA and mRNA levels produced on mvRNA templates NP71.11 and NP71.1. As shown in Fig. 4d-e, we found that although the replication levels remained consistent between the two polymerases, there was a significant increase in t-loop containing mvRNA transcription by the BM18 RNA polymerase relative to the WSN enzyme at 39°C.

To confirm that IAV transcription initiation is affected by the increase in temperature, we measured cap-snatching on model vRNA and cRNA promoters, as well as 71-nt long mvRNA derived from segments 5 (NP71.1). To this end, we first purified the WSN RNA polymerase and incubated it with a radiolabeled, capped 20-nt long RNA in the presence of model vRNA and cRNA promoters or NP71.1 in their positive or negative sense at 30°C, 37°C and 39°C. As shown in Fig. S8a, the endonuclease activity was minimal when no template was added to the reaction, while it was the most efficient when the vRNA promoter or a vRNA-sense mvRNA was present. The highest endonuclease activity was observed at 30°C. We also observed cap cleavage activity in the presence of the cRNA promoter and the positive-sense mvRNA templates. However, this activity was reduced relative to the negative sense templates at all temperatures. We next measured if transcription initiation was influenced by temperature. To this end, we incubated the purified IAV polymerase with a radiolabeled 11 nt long capped primer ending in 3’ AG or 3’ CA and either the wildtype vRNA promoter or mutant 3’ U1A promoter at 30°C, 37°C and 39°C (Fig. S8b). In this assay, the viral RNA polymerase aligns the capped primer to the terminal sequence of the template, and mismatched base-pair reduced the efficiency of transcription initiation. As shown in Fig. S8b, we observed a temperature-dependent reduction in transcription initiation but noted that initiation after base-pairing with nucleotide 2 of the primer was more temperature-independent in 2 of the 3 primer-template combinations. Together, these results are mostly in line with the reduced mRNA levels produced from the mvRNA templates we observe in our minigenome assays (Fig. 4). The results contrast the higher capped RNA levels produced from the full-length vRNA templates at 39°C during infection (Fig. 3), suggesting that the transcription efficiency of short mvRNA-like templates is different from NP-bound full-length RNA molecules. Presently, the underlying mechanism is not understood.

### Temperature dependent response to mvRNAs is RIG-I-dependent

To confirm that the activation of innate immune signaling by mvRNAs at 39°C was RIG-I-dependent, we utilized a HEK293 RIG-I -/- cell line. The knockout cells were transfected with mvRNA templates NP-71.11 and PB1-66, which both induce IFN-β promoter activity, and mvRNA templates NP-71.1 and PB1-62, which do not induce IFN-β promoter activity. In all transfections, we were unable to observe IFN-β promoter induction in the knockout cell line at 39°C, thus implying that the induction of IFN-β promoter activation by mvRNAs is RIG-I dependent at both 37°C and 39°C (Fig. S9b-c).

Finally, to confirm that the difference in IFN-β promoter activation at 39°C was not triggered by a higher activity of RIG-I, we performed an ATPase-hydrolysis assay using recombinant RIG-I. The ATPase activity of RIG-I was assessed in the presence of *in vitro* transcribed PA-66 based vRNA and cRNA at 37°C and 39°C. As negative controls, we also used alkaline phosphatase-treated vRNA and cRNA transcripts. As shown in Fig. 4f no significant difference was observed in the RIG-I activity at the two temperatures (Fig. 4f, Fig. S9d), which implies that the RNA binding-dependent ATP hydrolysis activity of RIG-I likely did not contribute to the observed increase IFN-β promoter activation at 39°C.

### Innate immune modulation by temperature is conserved across infections with H1N1 and H3N2 IAV strains

To confirm that temperature influences innate immune activation, we conducted infections using a panel of five additional influenza virus strains in HEK-luc cells: A/Melbourne/1946 (H1N1), A/Denver/1957 (H1N1), A/California/07/2009 (H1N1), A/Hong Kong/1968 (H3N2), and A/Brisbane/2007 (H3N2). All strains, with the exception of A/California/07/2009 (Cal09), exhibited a significantly elevated innate immune response at 39°C relative to 37°C, indicating a temperature-dependent enhancement of immune activation (Fig. 5a). The expression levels of viral proteins from Cal09 were markedly low, suggesting limited viral infection or replication capacity in our HEK-luc cells and explaining the minimal activation of the innate immune response observed.

**Fig. 5:**
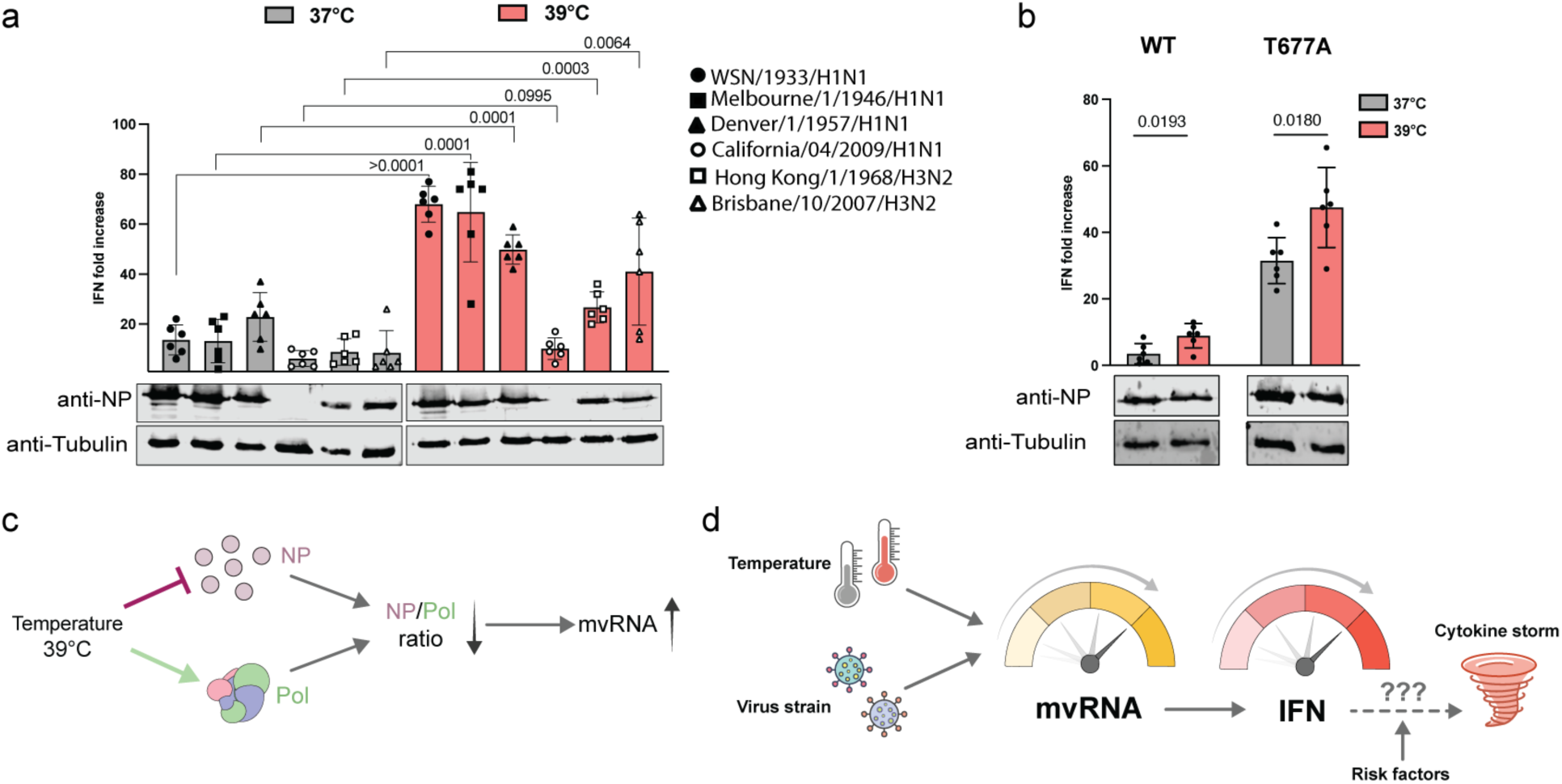
Temperature-driven modulation of innate immunity aligns with responses to other IAV strains. **a,** Analysis of IFN-β promoter activity, 12 hpi, in HEK-luc cell line adapted at 37°C and 39°C, infected with WSN, Melbourne46, Denver57, Cal09, HK68, and Brisbane07 IAV strains at an MOI of 3. The NP and tubulin expression was analyzed by western blot. **b,** IFN-β promoter activity was assessed in HEK-luc cells 12 hpi. The cells were adapted to 37°C and 39°C and infected at an MOI of 1 with either the WSN strain or the T677A mutant virus strain. The NP and tubulin expression was analyzed by western blot. Data are shown as the mean of 6 independent experiments. Error bars indicate standard deviation. *P* values were determined using a two-sided, unpaired *t*-test. **c,** An increase in infection temperature can differently impact polymerase and NP expression, and this imbalance may contribute to the synthesis of mvRNAs. **d**, Formation of mvRNAs is influenced by both temperature and the viral strain infecting the host, which can impact activation of the innate immune response, and cytokine and chemokine expression. A fever start may create a positive feedback loop that leads to increased mvRNA production and innate immune activation, and which may ultimately contribute to a cytokine storm. The likelihood for a cytokine storm may be modulated by risk and other host and viral factors.

Our model and previously published data indicate that both mvRNAs and ccRNAs are needed for innate immune activation, and that both are differentially expressed at 39°C. As a further validation for our model, we performed infections with a recombinant WSN virus containing a T677A substitution in PB1^16^. This mutation results in increased ccRNA levels and innate immune activation relative to wild-type WSN without increasing mvRNA levels. If our model were correct, elevated ccRNA starting levels would push the innate immune response further at 39°C. As shown in Fig. 5b, infection with WSN PB1 T677A led to higher ^16^elevated IFN-β reporter activity at 37°C relative to the wild-type strain, in line with our previous observations. Moreover, and as predicted, at 39°C the innate immune induction was further amplified, highlighting a temperature-dependent enhancement of the mutant’s immunostimulatory potential relative to the wild-type virus (Fig. 5c,d)

## Discussion

Despite its impact on human morbidity, the fever response remains one of the least understood aspects of an acute inflammatory reaction to infection. We here explored the impact of this increase in temperature on IAV non-canonical RNA synthesis and the innate immune response against IAV infection in vitro. We find that IAV RNA synthesis on full-length segments is not reduced at 39°C, but we do observe increased levels of segment 5 ccRNA synthesis and a corresponding decrease in NP levels. The reduced NP levels lead to an imbalance between the viral RNA polymerase and NP, which increases mvRNA synthesis and IFN-β promoter activation (Fig. 2 and 4). These dynamics can be captured by a relatively simple model of IAV infection (Fig. 1).

It is tempting to speculate that these dynamics could lead to a positive feedback loop in human or animal IAV infection. Following IAV infection and the initial production of mvRNAs, the innate immune response is activated and a moderate-grade fever triggered (e.g., 39°C). At this elevated temperature, the IAV RNA polymerase starts to produce additional mvRNAs, which sustain the activated innate immune and febrile response. It is possible that this prolonged response blocks the activation of negative feedback loops and/or contributes to a dysregulation of the innate immune response, which is a hallmark of infections with highly pathogenic influenza virus strains (Fig. 5c). This scenario would be particularly likely for infections with IAV strains that produce higher mvRNA base levels, such as the pandemic 1918 H1N1 IAV. At a higher base mvRNA level, the impact of fever would rapidly escalate mvRNA production and innate immune activation (Fig. 5d), potentially bringing the innate immune response to a dysregulated state.

Overall, our study incorporates cell culture-based assays, in vitro assays, and mathematical modeling, and offers a comprehensive dynamic perspective into the temperature-dependent effects on viral molecular machinery. In addition, the findings of this study underline the role of mvRNAs in innate immune activation and reveal that multiple rounds of aberrant RNA synthesis at febrile temperatures can stimulate innate immune activation.

## Material and methods

### Building a model for influenza replication with non-canonical RNA products

The model of the IAV replication dynamics was constructed by identifying the essential steps to capture replication and protein expression dynamics from a previously published model of intracellular IAV replication dynamics^23^. The steps include cRNA production from nuclear RNPs, vRNA production from cRNA, RNP assembly from polymerase, nucleoprotein, and c or vRNA, transcription of nuclear proteins making mRNA, translation of mRNA into protein, degradation of RNA, protein and RNP, and export of RNPs from the nucleus, gated by nuclear export protein. We then extend this model by three processes described in recent literature: i) Transcription of mRNA, influenza requires stealing CAP sequences produced by the host (CAP-snatching), ii) Replication requires a second polymerase complex to bind an RNP, and replication is only completed when the second complex binds to nucleoprotein, otherwise iii) incomplete replication leads to the formation of mvRNA. These products replicate to form positive and negative sense RNA products (mini vRNA or cRNA) and transcribe into capped aberrant mRNA products. On the other hand, we simplify the binding kinetics by assuming all complex formations follow the same diffusion-limited rate constant and are only distinguished by binding affinities (Table S1)^36^. From these assumptions, we formulate an ordinary differential equation model that describes the time evolution of the 23 species (Table S2). Except for CAP-snatching and RNP formation during replication, all processes are modeled using mass action kinetics. New RNP formation from replication follows a Michaelis-Menten like kinetics of the form:

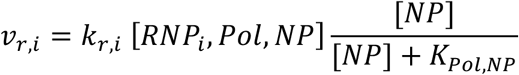

where *i* ∈ [*c*, *v*] denotes the RNA type, *k_r,i_*, is the respective replication rate, [*RNP_i_*, *Pol*, *NP*] is the number of c or vRNA RNP polymerase nucleoprotein complexes, [*NP*] is the number of nucleoproteins, and *K_NP,pol_* is the binding affinity of nucleoprotein to the polymerase complex. CAP-snatching kinetics are modeled similarly:

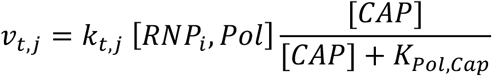

where *v_t,j_* with *j* ∈ [*np*, *pol*, *nep*] denotes the transcription rate for mRNA encoding nuclear protein, polymerase protein, and nuclear exit protein, respectively, [*RNP_i_*, *Pol*] is the number of c or vRNA RNP polymerase complexes, [CAP] is the size of the CAP-pool, and *K_pol,cap_*, is the binding affinity of the polymerase complexes to the CAP sequence.

The model was parameterized by directly or indirectly incorporating the parameter values identified in previous modeling efforts^23^ wherever possible. In the next step, all binding affinities were parameterized using data from *in vitro* assays. This left us with three undetermined parameters: The effective replication volume *V_r_*, the CAP synthesis rate *V_cap_*, and the CAP degradation rate *k_d,cap_* We found that volume *V_r_* needs to be a fraction (∼1%) of the nuclear volume for transcription to begin within 1-2 hours after infection. Considering that a single RNP only interacts with a small part of the nucleus, we compute an order of magnitude estimate of the CAP synthesis rate: *V_cap_* in the vicinity of the RNP is about one percent of the total mRNA synthesis rate, i.e, thousands of mRNA per minute. ^37^ ^38^

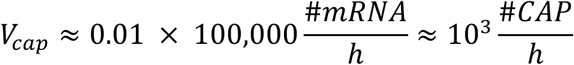

Finally, we set *k_d,cap_*, to match experimentally observed mRNA, cRNA, and vRNa dynamics^30^ (Fig. S1b-d). We recognized that the TPM values reported (transcript per kb per million) approximately match the RNA per cell quantified by RT-PCR^39^ or IAV infection of MDCK cells, and that RNA levels for infections in A549 cells are about 10-fold lower than in MDCK cells. We find *k_d,cap_*, = 0.1 by matching the 10x-scaled predictions for A549 cells (Fig S1a) to data measured for MDCK infection (Fig. S1b-d). We compute the time evolution of all species by numerically solving the ordering differential equations with initial conditions modeling the presence of a single viral particle and the steady state level of CAPs before infection, i.e. [*CAP*]*_t→∞_* = *V_cap_*/*k_d,cap_* To predict the effect on the innate immune response we assume that double stranded minis are the main driver and bind the RIG-I receptor with dissociation constant of 2.5 nM^24^.

### Cell Culture and Viruses

Human embryonic kidney cells (HEK293T) cells and alveolar basal epithelial cells (A549) cells were maintained in complete media (Dulbecco’s Modified Eagle’s Medium (DMEM, Sigma-Aldrich) supplemented with 10% fetal calf serum (FCS, Gibco) and 2 mM L-glutamine (Gibco)) at 5% CO2. HEK293T and Madin-Darby canine kidney cells (MDCK) were sourced from the American Type Culture Collection. Wild-type and *RIG-I* -/- human embryonic kidney cells stably expressing Firefly luciferase and GFP under the *IFN-b* promoter (HEK293-luc) were a generous gift from J. Rehwinkel (Oxford University) have been validated before^40^. Wild-type A549 were kindly provided by B. Ferguson (Cambridge University)^41^. A/WSN/1933 (H1N1) and T677AΔPB1 A/WSN/1933 (H1N1) virus were generated in the lab. Influenza A/Melbourne/1/1946 (H1N1), Influenza A/Denver/1/57 (H1N1), Influenza A/California/04/2009 (H1N1), Influenza A/Hong Kong/1/68 (H3N2), and Influenza A/Brisbane/10/2007 (H3N2) were ordered from BEI resources. Prior to all experiments, cells cultured at 37°C were split into independent, but paired flasks and incubated at 37°C and 39°C for 72 hours. Madin-Darby canine kidney cells used for plaque assays were incubated at 37°C.

### Plasmids

The pPolI plasmids encoding A/WSN/33 genome segments were previously described^42^. The pPolI plasmids encoding segment 2, 3- and segment 5-based templates with internal deletions were described in^14^ and Table S3; the pcDNA3 plasmids encoding wild-type A/WSN/33 RNP proteins, the plasmid expressing firefly luciferase under *IFN-b* promoter (pIFD(−116)lucter) and pcDNA3 encoding *Renilla* luciferase were described in ^14^.

### Virus infections

For infection experiments, A549 cells were seeded in 12-well plates at 2*10^5^ cells/well in complete DMEM. The next day, cells were infected with an MOI indicated for specific experiments in infection DMEM at 37°C, 5% CO_2_ for 1 h. After infection, cells were washed with PBS and incubated in infection DMEM at 37°C or 39°C, in 5% CO_2_. At the time periods indicated for specific experiments, supernatants were collected for virus titre quantification by plaque assay, and cells were lysed in either TRI Reagent (Sigma-Aldrich) for phenol-chloroform RNA extraction and isopropanol precipitation as described previously ^43^ or Laemmli buffer for western blotting.

### IAV growth kinetics at 37°C and 39°C

For single and multicycle IAV infections, A549 cells were seeded and incubated at the two temperatures. Cells were infected with A/WSN/33 at an MOI 3,1 or 0.01 PFU/cell in DMEM for 1 h. The inoculum was the removed, and cells were further incubated for different time points in growth media at the two temperatures, 37°C or 39°C. Supernatants were then titrated on MDCK cells by plaque assay at 37°C.

### RNP reconstitution and IFN-β promoter activity assay

HEK293T cells adapted at 37°C or 39°C for 3 days, were transfected in 12 or 24-well plate format with 0.25 μg/well of pcDNA3 plasmids encoding PB2, PB1, PA (wild-type WSN) and NP and 0.25 μg/well of pPolI plasmid encoding a full-length vRNA segment or an internally truncated segment (Table S3) under the control of the PolI promoter^14^. pCAGGS plasmids encoding PB2, PB1, PA and NP were used in case of 1918 H1N1 reconstitution assay. For assessment of *IFN-*β promoter activity, 0.1 μg of a plasmid expressing firefly luciferase from the *IFN-*β promoter and 0.01 μg of a plasmid expressing *Renilla* luciferase driven by the chicken β-actin promoter with a CMV enhancer were added to the transfection mix. The transfection was performed using a 1: 2.5 DNA (μg): Lipofectamine 2000 (Invitrogen) ratio. Twenty-four hours later, the DMEM was aspirated out. Next, we used PBS to detach the cells and split into 3 fractions: RNA isolation (1/2 cell pellet), western blotting (1/4 cell pellet) or the *IFN-*β promoter activity assay (1/4 cell pellet). For *IFN-*β promoter activity assays, 25 µl of DualGlo reagent (Promega) was added to 25 μl of cells suspended in PBS. This mixture was then incubated for 10 min and the firefly luciferase readings were taken using a Synergy LX plate reader (BioTek). Next, 25 µl of Stop-Glo reagent/well was added to each well, incubated for 10 min, and the *Renilla* luciferase readings were taken. For the final analysis, firefly luciferase values were normalized by the *Renilla* luciferase values.

### Cas13 assay

We used the previously established Cas 13 assay to quantify mvRNA in our sample^32^. The Cas13-based detection reactions contained 10 nM LbuCas13a, 20 mM HEPES pH 8.0, 60 mM KCl and 5% PEG, 2U/μL RNase inhibitor murine (New England Biolabs), 14 mM MgOAc, 0.25 μM 6UFAM (FAM-UUUUUUU-IowaBlack), 5 nM crRNA (GACCACCCCAAAAAUGAAGGGGACUAAAACAGAUAAUCACUCACAGAGUGACAU CGAA), and the reported amount of target RNA (approx. 50 ng/uL). Each reaction was first mixed in a volume of 44 μL in a 96-well plate. Following mixing, 20 μL of sample was transferred to 384-well plates in duplicate. The plate was then placed in a BioTek Synergy H1 Plate Reader and incubated at 37°C for 3 h. Fluorescence of each reaction was measured at 5 min intervals. The curves were analyzed as described previously^32^.

### mvRNA PCR

For the mvRNA RT-PCR in Fig. 1g, we used the method described previously^14^.

### Western blotting

For protein level analysis cells were first lysed in Laemmli buffer (62.5 mM Tris-HCl pH 6.8, 10% glycerol, 2% SDS, 100 mM DTT (Invitrogen), 0.01% Bromophenol Blue) followed by a short sonication. Next, proteins were separated using an 8% SDS-PAGE gel, followed by immobilization on a 0.45 µm or a 0.25 µm nitrocellulose membrane (GE healthcare). Next, the membranes were incubated in blocking buffer (PBS, 5% bovine serum albumin (RPI), 0.1% Tween-20 (RPI)) for 1 h. We then incubated the membranes with the primary antibody diluted in blocking buffer overnight at 4°C. Primary antibodies used in the study are described in Table S5. The following day, the membranes were washed 3x using PBST (PBS, 0.05% Tween-20). Binding of the primary antibodies was detected using secondary antibodies sourced from LI-COR (Table S5). After 2 h incubation with 1:10,000 dilution of secondary antibodies in blocking buffer at 4°C, membranes were washed 3x with PBST. Next, fluorescent signals were detected using an Odyssey DLx scanner (LI-COR). Image Studio Lite software (LI-COR) was used for quantification.

### Primer extension analysis

Primer extensions were performed essentially as described previously^43^. Briefly, total RNA was reverse-transcribed using Superscript III (Invitrogen) and ^32^P-labelled primers targeting the viral RNA of interest (vRNA, cRNA, mRNA) along with the cellular 5S rRNA loading control (Table S4) in 10 μl reactions at 50 °C for 1 h. Reactions were stopped using 10 μl Formamide loading dye (90% Formamide, 10 mM EDTA, 0.25% Bromophenol Blue, 0.25% xylene cyanol FF) and denatured at 95 °C for 2 min. cDNA products were resolved on a denaturing acrylamide gel (6% or 12% or 20% acrylamide (19:1, Bio-Rad)), 7 M urea, 1x Tris-borate EDTA buffer (2 mM EDTA, 89 mM boric acid, 89 mM Tris), 0.1% TEMED, 0.06% (w/v) APS) in 1x Tris-borate EDTA running buffer for 1.5-3 h at 35 W. In case of a 6% gel, the gel had to be dried. Finally, the radiolabeled signals were visualized using phosphor imaging plates (FujiFilm) on a Typhoon FLA 9000 scanner (GE Healthcare). Image Studio Lite software (LI-COR) was used for signal quantification. The quantified signal for RNA species was normalized to its corresponding 5S rRNA signal.

### IAV RNA polymerase purification

Wild-type IAV RNA polymerase was purified using tandem-affinity purification (TAP) as described previously ^43^ with modifications. Briefly, 4 μg of pcDNA3 plasmids expressing wild-type PB1, PA, and PB2-TAP were transfected into HEK293T cells using 60 µg PEI in Opti-MEM (Gibco). 48hours post transfection, cells were first harvested in PBS, followed by a wash with PBS. Cells were then lysed in 1 ml lysis buffer (50 mM HEPES pH 7.5, 200 mM NaCl, 25% glycerol (Sigma), 2% Tween-20 (RPI), 1 mM β-mercaptoethanol (Bio-Rad), and 1x EDTA-free Protease inhibitor cocktail (Roche)) at 4 °C for 1 h. The lysed cells were next sonicated to disrupt DNA and RNA molecules, and centrifuged at 17000 x *g* for 5 min at 4 °C. The cleared lysates were then bound to 50 μl IgG Sepharose beads 6 Flow (GE Healthcare). The beads were pre-washed 3x in binding buffer (50 mM HEPES pH 8, 200 mM NaCl, 25% glycerol, 2% Tween-20). The binding was performed under constant rotation at 4 °C for 2 hours. Next, the beads were washed 3x with binding buffer and 1x with cleavage buffer (50 mM HEPES pH 7.5, 200 mM NaCl, 25% glycerol, 0.5% Tween, 1 mM DTT). Finally, the TAP-tag was removed using Tobacco etch virus (TEV) protease (Invitrogen, #12575-015) in 250 μl of cleavage buffer. Cleavage was performed at 4°C for approximately 16 h. The beads were separated from the cleaved RNA polymerase by centrifugation at 500 *x g* for 1 min. The partially purified RNA polymerase was next analyzed through SDS-PAGE and silver staining was performed using a SilverXpress kit (Invitrogen).

### *In vitro* transcription and RNA transfection

DNA templates for *in vitro* transcription were prepared by PCR using the pPolI-PA66 plasmid as a template as described previously^14^. *In vitro* transcriptions were performed using MEGAshortscript™ T7 Transcription Kit (ThermoFisher) according to manufacturer’s protocol. The products were resolved using 15%denaturing PAGE (7M urea, 1x Tris-borate EDTA buffer, 0.1% TEMED, 0.06% (w/v) APS) in 1x Tris-borate EDTA running buffer. The products were gel-purified and desalted using an RNA Clean & Concentrator-5 (Zymo Research). The quality of the final RNA preparations was assessed using denaturing acrylamide gel electrophoresis.

### ATP hydrolysis assay

RIG-I was purified as described previously^44^. For ATPase assays, 0.5 µM RIG-I was incubated with 0.1 µM [γ-^32^P]ATP (6000 Ci/mmole, Perkin-Elmer) and 10 ng of in vitro transcribed template RNA as described above. Activity assays were performed in a buffer containing 50 mM HEPES pH 8.0, 150 mM NaCl, 2 mM MgCl_2_, and 5 mM DTT, and quenched using 1M formic acid/50 mM EDTA. [γ-^32^P]ATP and ^32^P_i_ were resolved using glass-backed PEI-cellulose TLC plates (Sigma-Aldrich) in 0.4 M KH_2_PO_4_ (pH 3.4). After wrapping the TLC plates in plastic foil, the radioactive signals were detected using BAS-MS phosphor imaging plates (FujiFilm) through a 1 h exposure and visualized using a Typhoon FLA 9000 scanner (GE Healthcare). Densitometry analysis was performed using ImageJ.

### In-vitro activity/transcription assays with the purified IAV polymerase

To measure the ability of the IAV RNA polymerase to extend a capped primer, first a synthetic 11-nt long RNA with 5ʹ diphosphate (Chemgenes; ppGAAUACUCAAG) was capped with a radiolabeled cap-1 structure in 20 µl reactions containing 1 µM RNA, 0.25 µM [α-^32^P]GTP (3000 Ci/mmole, Perkin-Elmer), 0.8 mM S-adenosylmethionine, 0.5 U/µl *Vaccinia virus* capping enzyme (NEB), and 2.5 U/µl 2ʹ-O-methyltransferase (NEB) at 37°C for 1 h. The capped RNA was purified using an oligonucleotide cleanup kit (Zymo Research) and eluted in 30 µl water. To test the transcriptional activity of the IAV RNA polymerase, we used 4-μl reactions containing 5 mM MgCl_2_, 1 mM DTT, 2 U/μl RNase inhibitor (APExBIO), 0.5 μM of RNA template (Table S3), 500 μM ATP, 500 μM CTP, 500 μM GTP, 0.2 µl capped RNA primer, 12.5% glycerol, 1% Tween-20, 100 mM NaCl, 25 mM HEPES pH 8, and 5 ng/µl RNA polymerase. Reactions were incubated at 30°C, 37°C, and 39°C for 30 minutes and analyzed with 12% denaturing PAGE and autoradiography. To measure the ability of the IAV RNA polymerase to cleave a capped primer, a synthetic 20-nt long RNA with 5ʹ diphosphate (Chemgenes; pp AAUCUAUAAUAGCAUUAUCC) was first capped with a radiolabeled cap-1 structure as described above for the 11-nt long primer. To test the cleavage activity, 4-µl reactions were performed as described above, but NTPs were omitted and replaced with RNase-free water.

### RNA isolation

Total RNA was extracted from cells using trizol, chloroform method as previously described ^43^. RNA concentration was assessed using the Qubit 2 fluorometer (Thermo Fisher Scientific, USA) with Qubit RNA HS Assay Kit (Thermo Fisher Scientific, USA). The quality of total RNA expressed as RNA Integrity Number (RIN) was determined with Bioanalyzer 2100 instrument (Agilent, USA) using an Agilent RNA Pico 6000 Kit (Agilent, USA). The threshold RIN reading greater than 8.0 was taken as cut-off point for transition to the stage of library preparation.

### Library preparation and sequencing

A total of 24 cDNA libraries were prepared from three biological replicates of 12 h mock and infected A549 and HEK293T cell line respectively. Purified total RNA samples were first examined on Bioanalyzer 2100 using RNA 6000 Pico chip to evaluate the integrity and concentration (Agilent Technologies, CA), then normalized onto 96-well plates. The poly-A containing RNA transcripts in these samples were converted to cDNA using barcoded oligo-dT primers in the reverse transcription reaction to index each sample following the drop-seq method ^45^. The pooled barcoded cDNA samples were amplified by PCR and purified, then turned into sequencing libraries using the Illumina Tagment DNA Enzyme and Buffer kit (Illumina, CA) to include only the poly-A tail adjacent 3’ ends of RNA transcripts. These libraries were examined on the Bioanalyzer (Agilent, CA) DNA HS chips for size distribution and quantified by Qubit fluorometer (Invitrogen, CA), then sequenced on Illumina NovaSeq 6000 S Prime flowcell using the 100 cycle v1.5 kit. Raw sequencing reads were filtered by Illumina NovaSeq Control Software and only the Pass-Filter (PF) reads were used for further analysis.

### RNA seq analysis

The raw data were saved as FASTQ format files. The quality control of the raw and trimmed reads was performed using fastp^46^.The reads complementary to the genome of influenza A/WSN/1933 (H1N1) were filtered out from the trimmed reads and the filtered reads were used for transcript quantification using RNA STAR tool (GRCh38_RefSeq_Transcripts)^47^. edgeR was used to convert the transcript quantifications to gene quantifications^48^.

### Differential gene expression analysis

To study genes involved in the cellular response to influenza A virus infection in A549 cells adapted to 37 or 39°C and mock infected HEK293T cells, differentially expressed genes between the temperatures were identified following a standard workflow using edgeR with an FDR-adjusted *p*-value < 0.05 and the absolute value of a log2(FC) > 1)^48^. Reactome pathway enrichment analysis was used to categorize the top hits from the differentially expressed genes using R package ReactomePA^49^.

### TSO-based RT-PCR

For the TSO-based RT-PCR, RNA was collected from A549 cells infected with an MOI of 1 of A/WSN/33 virus or mock-infected. After extraction, 1 µg was either treated or mock-treated with oligo d(T)_25_ beads from the magnetic mRNA isolation kit (NEB) according to manufacturer’s instructions and purified using the RNA Clean & Concentrator-5 kit (Zymo Research). Equal volumes (containing around 100 ng) of RNA were used for TSO-based RT. To this end, RNA was first denatured in the presence of dNTPs (1mM final concentration) and either the Tuni-13 LNA3 (ACGCGTGATCAGTAGAAA+CA+AG+G) or the oligo d(T)_20_ primer (1 µM final concentration) in 3 µl volume at 70 °C for 5 min. Denatured RNAs were immediately placed on ice for 5 min. Next, 2 µl of enzyme mix containing template switching RT buffer (NEB), template switching RT enzyme mix (NEB),and the TSO (GCTAATCATTGCAAGCAGTGGTATCAACGCAGAGTACATrGrGrG, 3.75 µM final concentration) were added to the RNA mix and incubated at 42 °C for 90 min. Reactions were terminated by incubation at 85 °C for 5 min and subsequently cooled to 4 °C. The unused primers were digested with Thermolabile Exonuclease I (NEB) and reactions were diluted 2x with water. One µl of diluted RT reaction was used for PCR amplification with Q5 High-Fidelity DNA polymerase (NEB) using TSO-specific forward primer (CATTGCAAGCAGTGGTATCAAC) and either an NP-specific (GATTTCGATGTCACTCTGTGAG) or a PA-specific (GGATTGAAGCATTGTCGCAC) reverse primer. Thermocycling conditions included: i) initial denaturation at 98 °C for 30 s; ii) 20-30 cycles of denaturation at 98 °C for 10 s, annealing at 50 °C for 15 s and extension at 72 °C for 10 s; iii) final extension at 72 °C for 2 min. Note, that different numbers of cycles were used for 2-4h and 6-10h time points to avoid oversaturation. The PCR products were resolved on a 10% TBE acrylamide gel and visualized using SYBR Gold (Invitrogen).

### Statistical analysis

All data were represented as mean ± SD. Data were analyzed in GraphPad Prism 10.0 software using Student’s t-test. Statistical significance was set at a threshold of P < 0.05. Data are representative of n = 3 biological replicates, unless stated otherwise in the figure legend.

## Acknowledgements

The authors would like to thank Shirley Yang for sharing work on a related research project, and Wei Wang, Jennifer Miller and Jean Arly Volmar in the Princeton Genomics Core Facility for assistance with the RNAseq.

## Author contributions

KB and AJWtV conceived hypotheses. KB created constructs and performed infection and transfection experiments. KB and AJWtV analyzed biological data. DRW developed the model to capture IAV infection cycle and aberrant RNA synthesis with input from KB and AJWtV. CHL performed Cas13 assay. EE performed TSO assay. AJWtV and KB drafted the manuscript. All authors commented on and approved the manuscript.

## Data availability

All data needed to evaluate the conclusions are available in the paper and Supplemental Materials. The raw RNA-seq data sets generated during this study are available through NCBI’s BioProject database under accession number PRJNA1174240. Additional raw data are available via 10.5281/zenodo.15213082. Plasmids are available from the authors or from Addgene.

## Code availability

All code used to simulate IAV infection dynamics are available at GitHub https://github.com/weilandtd/Influenza-infection-cycle.

