## Supplemental Figures for "Febrile temperature activates the innate immune response by promoting aberrant influenza A virus RNA synthesis"

### Febrile temperature promotes aberrant influenza A virus RNA synthesis and triggers innate immune responses

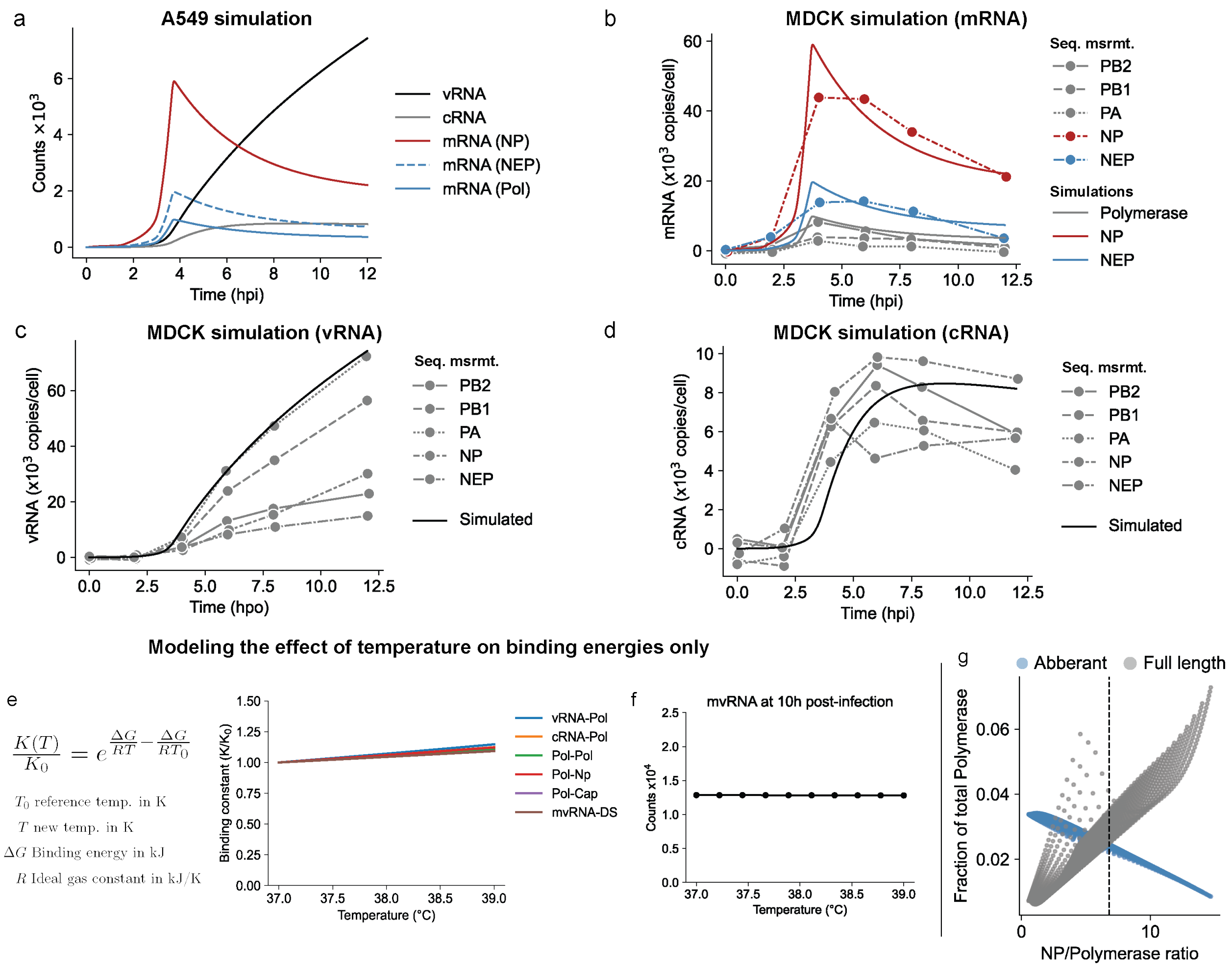


**Supplementary Fig.1**| **a,** Model prediction of cRNA, vRNA and mRNA subspecies for single infection cycle of IAV in A549 cell line. Note that we assume that the replication of the eight IAV segments (vRNA and cRNA) is not fundamentally different, and that the expression differences among the segments are driven by variations in transcription (mRNA) efficiency, which is why only one vRNA and cRNA are simulated here. Comparing the model predictions to experimental data collected in MDCK cells for **b,** mRNA levels for polymerase segments, nucleoprotein and nuclear exit protein, **c,** vRNA level, **d,** cRNA level. Model predictions for temperature effects on only binding constant **e,** Mathematical expression and simulation for temperature variation on binding constants. **f,** mvRNA level as function of the temperature effect on binding constants only. **g,** Fraction of polymerase engaged in transcription of aberrant and full-length products as a function of NP/polymerase ratio.

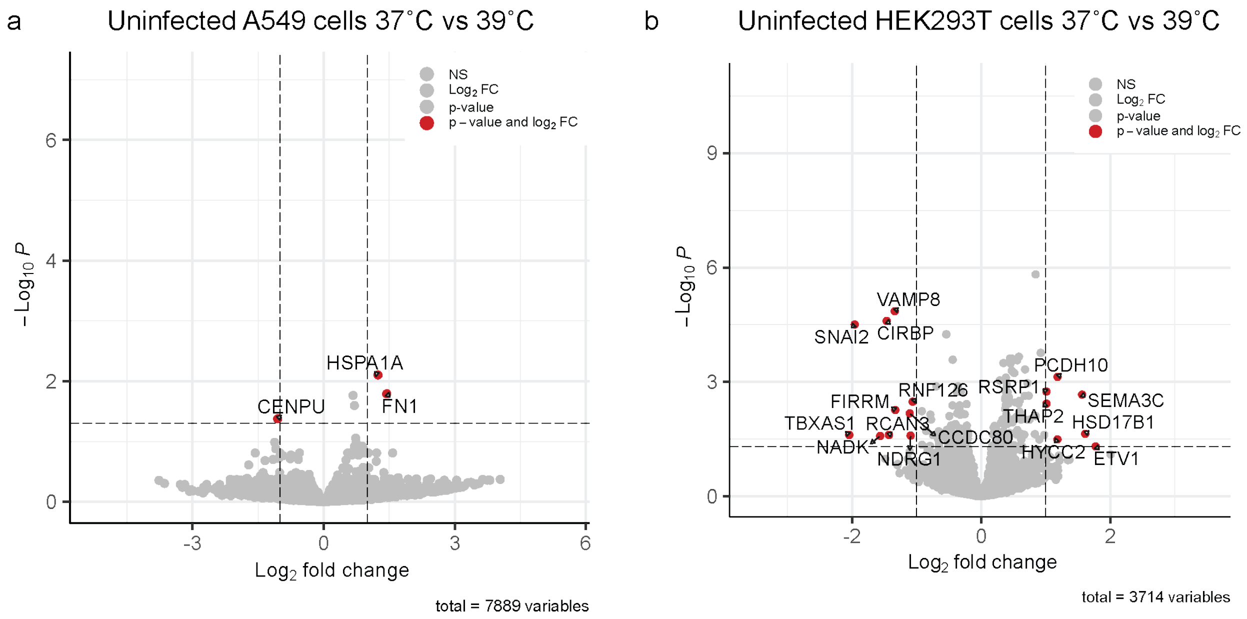


**Supplementary Fig.2**| **a,** Volcano plot illustrating differentially expressed genes in A549 mock infected cells after being adapted to 37˚C and 39˚C for 3 days **b,** Volcano plot illustrating the differentially expressed genes in HEK293T cells after being adapted to 37˚C and 39˚C for 3 days. RNA-seq was performed on 3 biological replicates.

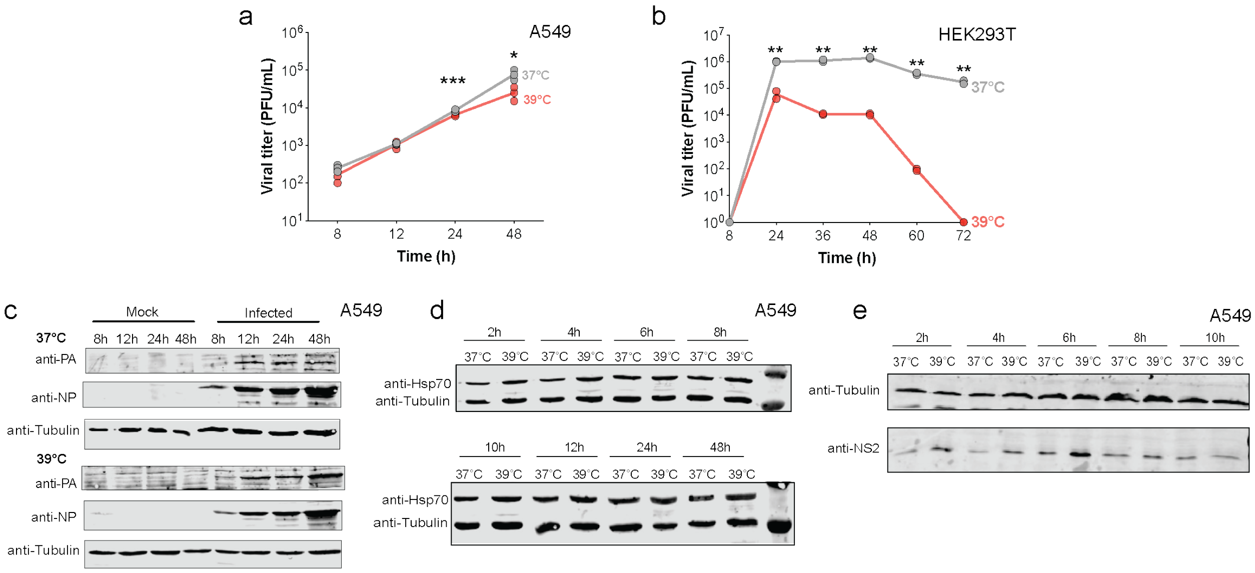


**Supplementary Fig.3**| **a.** Measurement of steady state vRNA levels for full-length PA at 8 hours post-infection (hpi) and 12 hpi in A549 cells infected with WSN virus, with cells adapted to 37°C and 39°C. Growth kinetics of lab adapted WSN virus in A549 (**a**) and HEK293T (**b**) cells at different temperatures (37°C, or 39°C). A549 cells or HEK293T cells were infected at an MOI of 0.01 at different temperatures 37°C and 39°C. The supernatants of the infected cells were harvested at the indicated times, and the virus titers were determined by performing plaque assays in MDCK cells at 37°C. Data are shown as the mean of triplicate experiments. Error bars indicate the standard deviation. The *P* values were determined by using an unpaired *t* test. (**P* < 0.05; ***P* < 0.01; ****P* < 0.001). **c,d,e,** Cell lysates were collected at different time points from infected and mock cells and analyzed by immunoblotting with indicated antibodies.

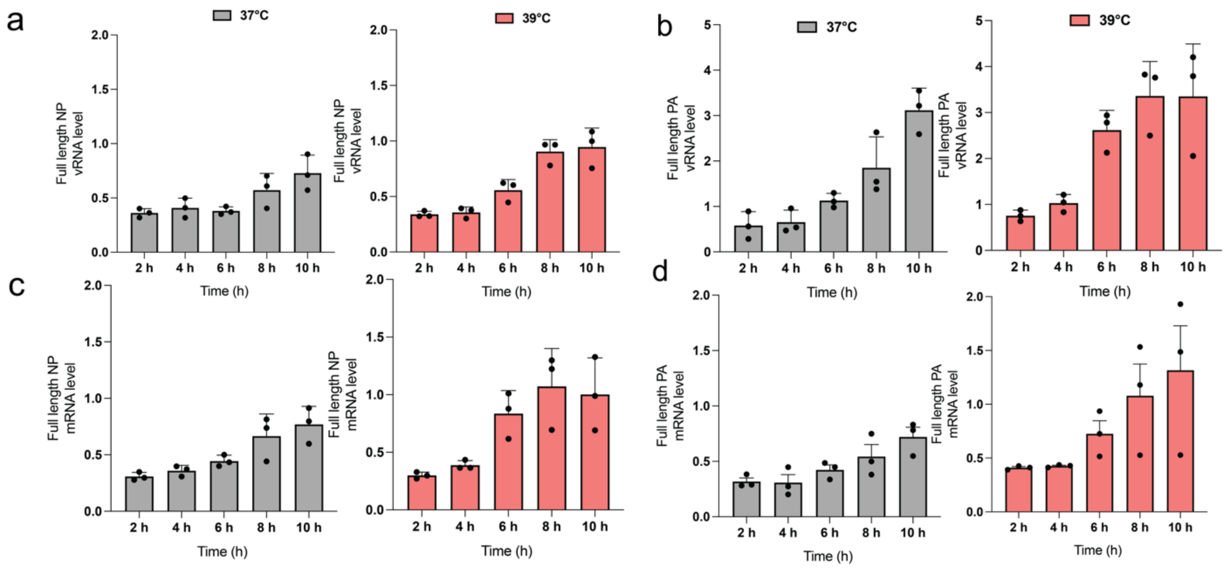


**Supplementary Fig.4**| A549 cells adapted at 37˚C and 39˚C were infected with an MOI of 1 of A/WSN/33 and RNA samples were taken at different time points post-infection. **a,b**, Quantification of steady state vRNA levels and **c,d,** mRNA levels for NP and PA segments, respectively.

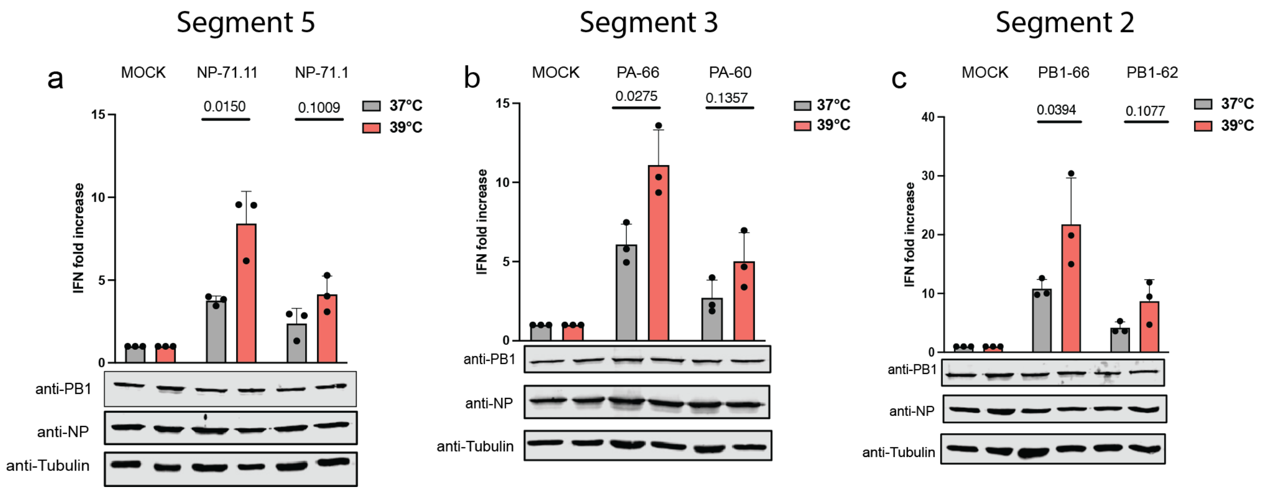


**Supplementary Fig.5**| **a,b,c,** Analysis of IFN-β promoter activity induced by the replication of segment 5, 3 or 2 mvRNAs by the WSN RNA polymerase. PB1, NP and tubulin expression was analyzed by western blot. Data are shown as the mean of triplicate experiments. Error bars indicate the standard deviation. The *P* values were determined by using an unpaired *t* test.

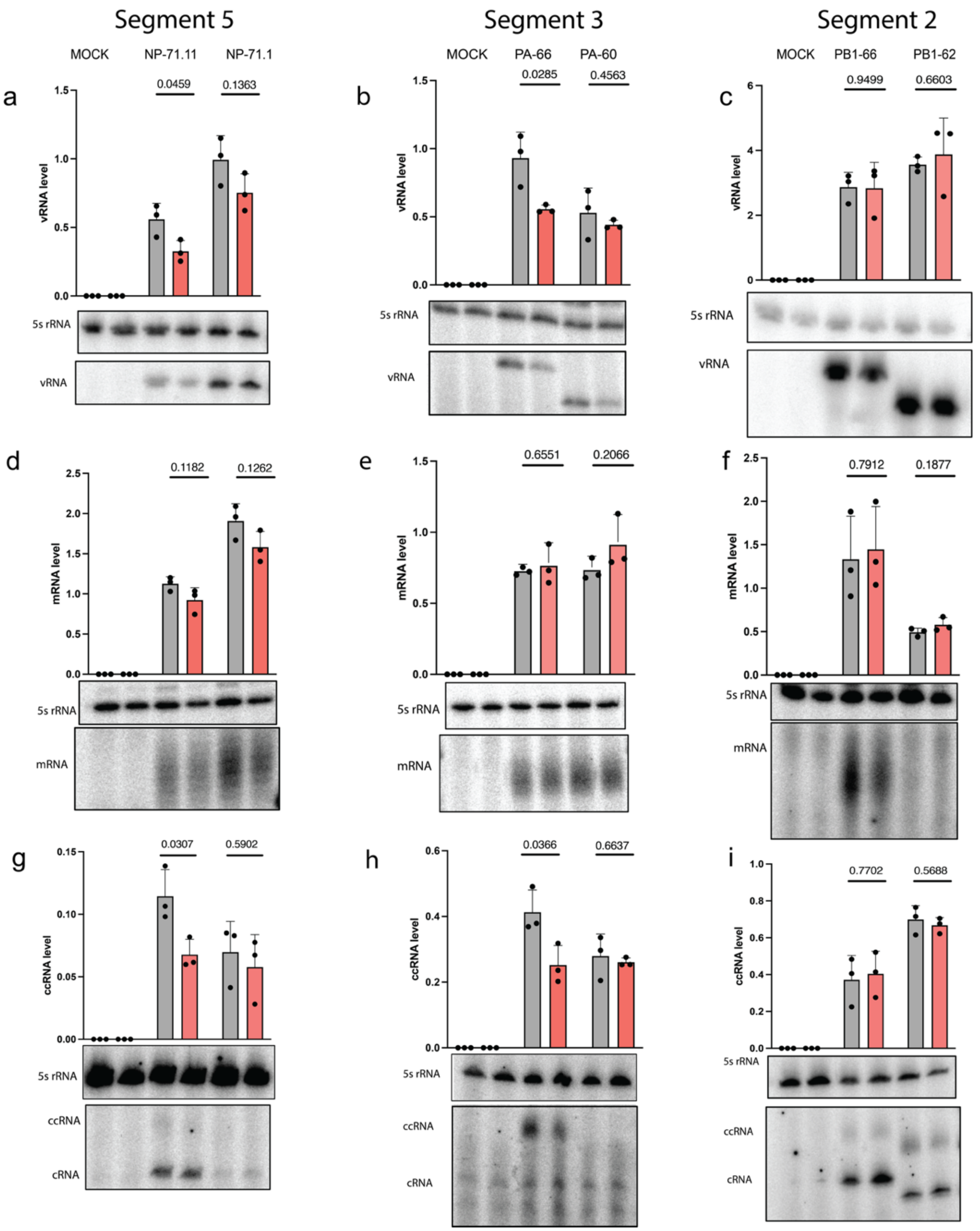


**Supplementary Fig.6**| **a,b,c,** Steady state mvRNA and 5S rRNA levels quantified by primer extension in the presence 24 hours post-transfection of BM18 IAV RNA polymerase. **d,e,f**, Steady state mRNA and 5S rRNA levels analyzed 24 hours post-transfection of BM18 IAV RNA polymerase. **g,h,i**, Steady state cRNA and ccRNA levels analyzed 24 hours post-transfection of BM18 IAV RNA polymerase. Data are shown as the mean of 3 independent experiments. Error bars indicate standard deviation. *P* values were determined using a two-sided, unpaired *t*-test.

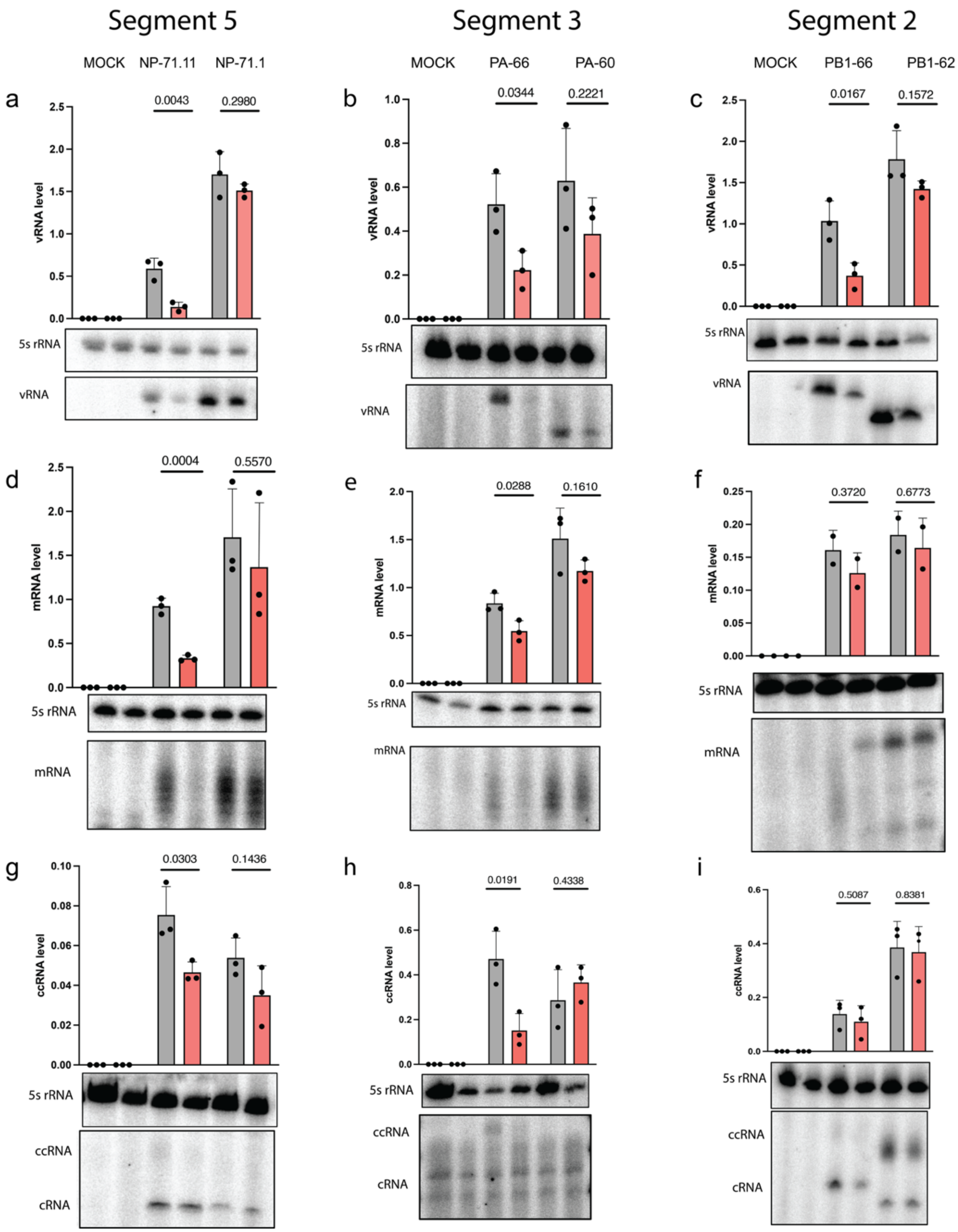


**Supplementary Fig.7| a,b,c,** Steady state mvRNA and 5S rRNA levels measured and quantified, 24 hours post-transfection by primer extension in the presence of WSN RNA polymerase. **d,e,f**, Steady state mRNA and 5S rRNA levels measured and quantified, 24 hours post-transfection by primer extension in the presence of WSN RNA polymerase. **g,h,i**, Steady state cRNA and ccRNA levels measured and quantified, 24 hours post-transfection by primer extension in the presence of WSN RNA polymerase. Data are shown as the mean of triplicate experiments. Error bars indicate the standard deviation. The *P* values were determined by using an unpaired *t*-test.

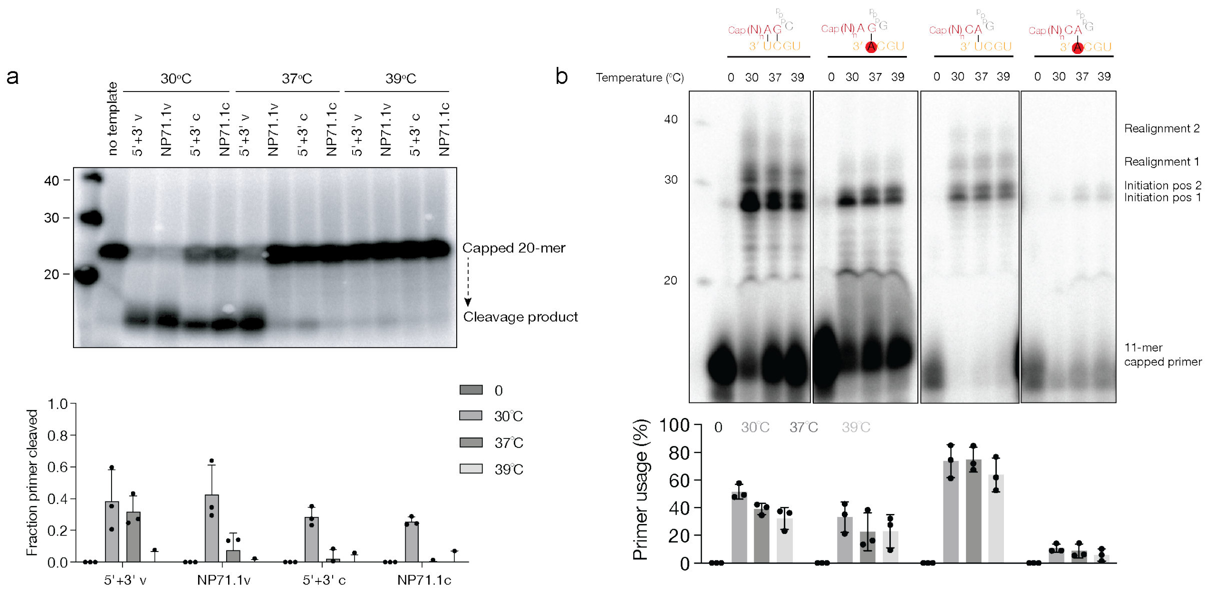


**Supplementary Fig.8| a,** Cap-snatching of a $^{32}$P-labeled capped, 20-nt long RNA primer by the IAV RNA polymerase at 30˚C, 37˚C and 39˚C. Cleavage reactions were analyzed by denaturing PAGE. Lower panel shows fraction of the primer cleaved at three temperatures. **b,** Extension of a radiolabeled, capped, 11-nucleotide-long RNA primer ending in 3′ AG on the wild-type or 3′ 1U→A vRNA promoter and 3′ CA on the wild-type or 3′ 1U→A vRNA promoter, at 30˚C, 37˚C and 39˚C. Quantitation of data in the lower panel shows percentage of primer used at the three temperatures.

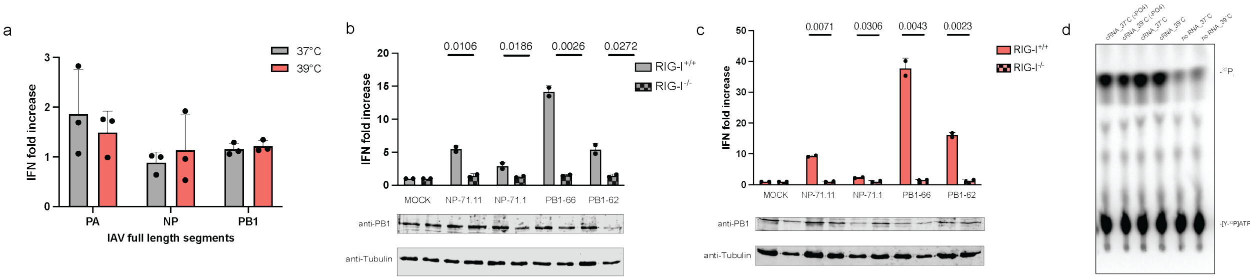


**Supplementary Fig.9| a,** IFN-$\beta$ promoter activity induced by the expression of full-length segment 2, 3 and 5 in HEK293T cells adapted at 37˚C and 39˚C. **b,** IFN-$\beta$ promoter activity induced by expression of segment 2 and 5 mvRNAs in in wild-type (RIG-I$^{+/+}$) or RIG-I knockout (RIG-I$^{-/-}$) HEK293 cells at the two temperatures. Bottom two panels show western blot analysis**.** Data are shown as the mean of two independent experiments **c,** ATPase activity of recombinant RIG-I was assessed in the presence of *in vitro* transcribed PA66-based cRNA, and dephosphorylated cRNA, as well as no RNA control. Data are shown as the mean of triplicate experiments unless specified. Error bars indicate the standard deviation. The *P* values were determined by using an unpaired *t* test.

**Table S1: Model parameters at 37 and 39˚C.**

| Symbol | Description | Source | Value 37˚C | Value 39˚C | Unit |
|---|---|---|---|---|---|
| $k$ | Universal diffusion limited binding constant | 36 | $2.78 \times 10^5$ | $2.78 \times 10^5$ | 1/h/M |
| $K_{vRNA/Pol}$ | Binding constant of vRNA to polymerase complex | 22 | 0.5 | 0.57 | nM |
| $K_{cRNA,Pol}$ | Binding constant of cRNA to polymerase complex | 23 | 13 | 14.6 | nM |
| $K_{Pol,Pol}$ | Binding constant two polymerase complexes | assumption | 100 | 101 | nM |
| $K_{Pol,NP}$ | Binding constant polymerase to first nucleoprotein | 24,25,26 | 15 | 16.8 | nM |
| $K_{Pol,Cap}$ | Binding constant polymerase to host mRNA cap | 27 | 1 | 1.1 | µM |
| $V_{cap}$ | Synthesis rate for caps | methods | $10^3$ | $10^3$ | #CAP/h |
| $k_{d,cap}$ | Degradation rate for caps | assumption | 0.1 | 0.1 | 1/h |
| $k_{t,NEP}$ | Transcription rate of nuclear export protein | 21* | 35 | 14 | 1/h |
| $k_{t,Pol}$ | Transcription rate of polymerase mRNA | 21* | 17.5 | 14 | 1/h |
| $k_{t,NP}$ | Transcription rate of nucleoprotein | 21* | 105 | 42 | 1/h |
| $k_{t,mini}$ | Transcription rate of mvRNA | 21* | 3.5 | 3.5 | 1/h |
| $k_{tl}$ | Universal translation rate | 21 | 6 | 6 | 1/h |
| $k_{rv}$ | Replication rate of vRNA (from cRNA) | 21 | 13.85 | 13.85 | 1/h |
| $k_{rc}$ | Replication rate of cRNA (from vRNA) | 21 | 1.38 | 1.38 | 1/h |
| $k_{r,mini}$ | Initiation rate of mvRNA rate (from vRNA) | assumption | 1.38 | 1.38 | 1/h |
| $k_d$ | Degradation rate of mRNA | 21 | 0.33 | 0.33 | 1/h |
| $k_{d,vc}$ | Degradation rate of nascent cRNA and vRNA | 21 | 36 | 36 | 1/h |
| $k_{d,RNP}$ | Degradation rate of RNPs | 21 | 0.09 | 0.09 | 1/h |
| $k_{d,p}$ | Degradation rate of proteins | 21 | 0.5 | 0.5 | 1/h |
| $k_{exp}$ | Export rate of RNPs per NEP | 21 | $10^{-8}$ | $10^{-8}$ | 1/h/#NEP |
| $z$ | Stoichiometry of nucleoprotein per RNP | 12 | 20 | 20 | #NP/#POL |
| $V_r$ | Viral replication volume in nucleus | methods | $10^{-15}$ | $10^{-15}$ | L |

*Computed from the transcription rate per nucleotide reported.

**Table S2: Model species**

| Symbol | Description |
|---|---|
| $[CAP]$ | Pools of CAP sequences in the vicinity of the RNP |
| $[S_v]$ | Representative RNP with viral RNA* |
| $[S_c]$ | Representative RNP with complementary RNA* |
| $[S_v]_{cyt}$ | Exported representative RNP with viral RNA* (cytosolic) |
| $[NP]$ | Nucleoproteins |
| $[NEP]$ | Nuclear exit protein |
| $[Pol]$ | Polymerase protein** |
| $[mRNA_{NP}]$ | mRNA encoding nucleoproteins |
| $[mRNA_{NEP}]$ | mRNA encoding nuclear export protein |
| $[mRNA_{Pol}]$ | mRNA encoding RNA polymerase protein subunit** |
| $[S_v, Pol]$ | Representative vRNP polymerase complex*,** |
| $[S_c, Pol]$ | Representative cRNP polymerase complex*,** |
| $[S_v, Pol, NP]$ | Representative vRNP polymerase complex*,** |
| $[S_c, Pol, NP]$ | Representative cRNP polymerase complex*,** |
| $[mcRNA]$ | Mini viral RNA positive sense |
| $[mvRNA]$ | Mini viral RNA negative sense |
| $[mcRNA, Pol]$ | Mini viral RNA positive sense polymerase complex** |
| $[mvRNA, Pol]$ | Mini viral RNA negative sense polymerase complex** |
| $[mcRNA, Pol, Pol]$ | Mini viral RNA positive sense two polymerase complex** |
| $[mvRNA, Pol, Pol]$ | Mini viral RNA negative sense two polymerase complex** |
| $[cap.mcRNA]$ | Capped mini viral RNA positive sense |
| $[cap.mvRNA]$ | Capped mini viral RNA negative sense |
| $[ds.mvRNA]$ | Double stranded mini viral RNA |

* Representative RNP meaning average across all 8 segments

** Polymerase protein, meaning sum across the 3 polymerase subunits

**Table S3**

| Template Name | Sequence of the RNA templates (5′ to 3′) |
|---|---|
| NP STOP | WSN NP full-length STOP (Start codon ATG mutated in stop codon TCG) |
| PA | WSN PA full length |
| PB1 STOP | WSN PB1 full length STOP (Start codon ATG mutated in stop codon TCG) |
| NP246 | AGUAGAAACAAGGGUAUUUUUCUUUAAUUGUCGUACUCCUCUGCAUUGUCUCCGAAG AAAUAAGAUCCUUCAUUACUCAUGUCAAAGGAGGGCACGAUCGGGCUCGUUGCCUUU UCGUCCGAGAGCUCGAAGACUCCCCGCCCCUGGAAAGACACUAGUCUCCAUCUGUUC GUAAGAUCGUUUGGUGCCUUUGGUCGCCAUGAUUUCGAUGUCACUCUGUACUAGUC UACCCUGCUUUUGCU |
| NP71.1 | AGUAGAAACAAGGGUAUUUUUCUUUACUAGUUAGGUAGUAUACCUAGUAACUAGUCU ACCCUGCUUUUGCU |
| NP71.11 | AGUAGAAACAAGGGUAUUUUUCUUUACUAGUGGCAGCAAAAGCACCCAUACUAGUCU ACCCUGCUUUUGCU |
| PA-60 | AGUAGAAACAAGGUACUUUUUUGGACAGUAUGCCAUUUUGAAUCAGUACCUGCUUUC GCU |
| PA-66 | AGUAGAAACAAGGUACUUUUUUGGACAGUAUGGAUAGCACAUUUUGAAUCAGUACCU GCUUUCGCU |
| PB1-66 | AGUAGAAACAAGGCAUUUUUUCAUGAAGGACAAGCUAAACAUUCAAAUGGUUUGCCU GCUUUCGCU |
| PB1-62 | AGUAGAAACAAGGCAUUUUAAGUCGGAUUGACAUCCAUUCAAAUGGUUUGCCUGCUU UCGCU |
| 5S 100 | TCCCAGGCGGTCTCCCATCC |
| cRNA 3’ | GGCCUUGUUUCUACU |
| cRNA 5’ | AGCGAAAGCAGGCC |
| vRNA 3’ | GGCCUGCUUUUGCU |
| vRNA 5’ | AGUAGUAACAAGGCC |

**Table S4: DNA oligonucleotides used for primer extension.**

| Primer name | Target RNA | DNA oligonucleotide (5’ to 3’) |
|---|---|---|
| NP- | Non-full-length NP vRNA and aberrant products | AGCAAAAGCAGGGTAGACTAGT |
| NP+ | Non-full-length NP mRNA | ACTAGTCTACCCTGCTTTTGC |
| NP5’ | Non-full-length NP cRNA | AGTAGAAAACAAGGGTATTTTTC |
| PA_PEplus2 | Non-full-length PA cRNA | AGTAGAAACAAGGTACTTTTTTGGACA |
| PA vRNA | Non-full-length PA vRNA and aberrant products | AGCGAAAGCAGGTACTGATTC |
| PA_PEplus_short | Non-full-length PA mRNA | TTTTGAATCAGTACCTGCTTTCG |
| PB1-3’ | Non-full-length PB1 vRNA | AGCGAAAGCAGGCAAACCATTTG |
| PB1 cRNA | Non-full-length PB1 cRNA and aberrant products | AGTAGAAACAAGGCATTTT |
| PB1_PEplus_short | Non-full-length PB1 mRNA | CAAATGGTTTGCCTGCTTTCG |
| PA-2119 | Full length PA vRNA | GGGTTTTGCCTGCTTTCG |
| PA-135 | Full length PA cRNA and mRNA | TGCTGCAAATTTGTTTGTTTCG |
| NP149- | Full length NP vRNA | ATTTCTTCGGAGACAATGCAG |
| NP149+ | Full length NP cRNA and mRNA | TAAGATCGTTTGGTGCCTTTG |
| 5S_100 | 5S sRNA; loading control | TCCCAGGCGGTCTCCCATCC |

**Table S5: Antibody used for western blot.**

| | | |
|---|---|---|
| Primary antibody | | |
| NP | Rabbit, GTX125989, GeneTex | 1:2000 |
| PB1 | Rabbit, GTX125923, GeneTex | 1:1000 |
| PB2 | Rabbit, GTX125926, GeneTex | 1:1000 |
| PA | Rabbit, GTX125932, GeneTex | 1:1000 |
| Hsp70 | Rabbit, 24532, Cayman Chemicals | 1:500 |
| NS1 | Rabbit, PA5-32243, Invitrogen | 1:1000 |
| $\gamma$-tubulin | Rat, MCA77G, Bio-Rad | 1:5000 |
| Secondary antibody | | |
| IRDye 680 goat anti-rat | 926-68076, LI-COR | 1:10000 |
| IRDye 800 donkey anti-rabbit | 926-32213, LI-COR | 1:10000 |
